# LipiGo: A Versatile DNA-Lipid Nanoparticle Hybrid for Precision Drug Delivery

**DOI:** 10.1101/2025.09.15.676334

**Authors:** Karoline Kadletz, Ceren Kimna, Valentin Kenneth Reichenbach, Olga Carofiglio, Denise Jeridi, Igor Khalin, Ying Chen, Izabela Horvath, David-Paul Minde, Mayar Ali, Taras Sych, Sofia Hu, Luciano Hoeher, Eren Aydeniz, Yagmur Ayse Kulcu, Jakub Grzejdak, Alessio Ricci, Erdinc Sezgin, Nikolaus Plesnila, Arthur Liesz, Siegfried Ussar, Farida Hellal, Markus Elsner, Ali Ertürk

## Abstract

Lipid nanoparticles (LNPs) are established carriers for nucleic acid delivery, however, achieving efficient delivery to non-hepatic tissues remains a major challenge. Here, we present LipiGo, a nanocarrier platform engineered by integrating short single stranded DNA molecules into the lipid nanoparticle structure. Using whole body tissue clearing, advanced imaging and AI-based analysis, we show that LipiGo redirects functional mRNA delivery to lymphoid organs, particularly the spleen. Immune cell profiling further reveals enhanced uptake within key immune populations, including antigen-presenting cells, compared to standard LNPs. Beyond passive redistribution, LipiGo leverages DNA hybridization to enable modular attachment of targeting ligands for active targeting as demonstrated by cell-specific delivery to white adipocytes. Overall, the dual-purpose design principle of LipiGo demonstrates high modularity and efficiency, enabling tissue and cell specific delivery beyond hepatic applications.

**Graphical Abstract:** 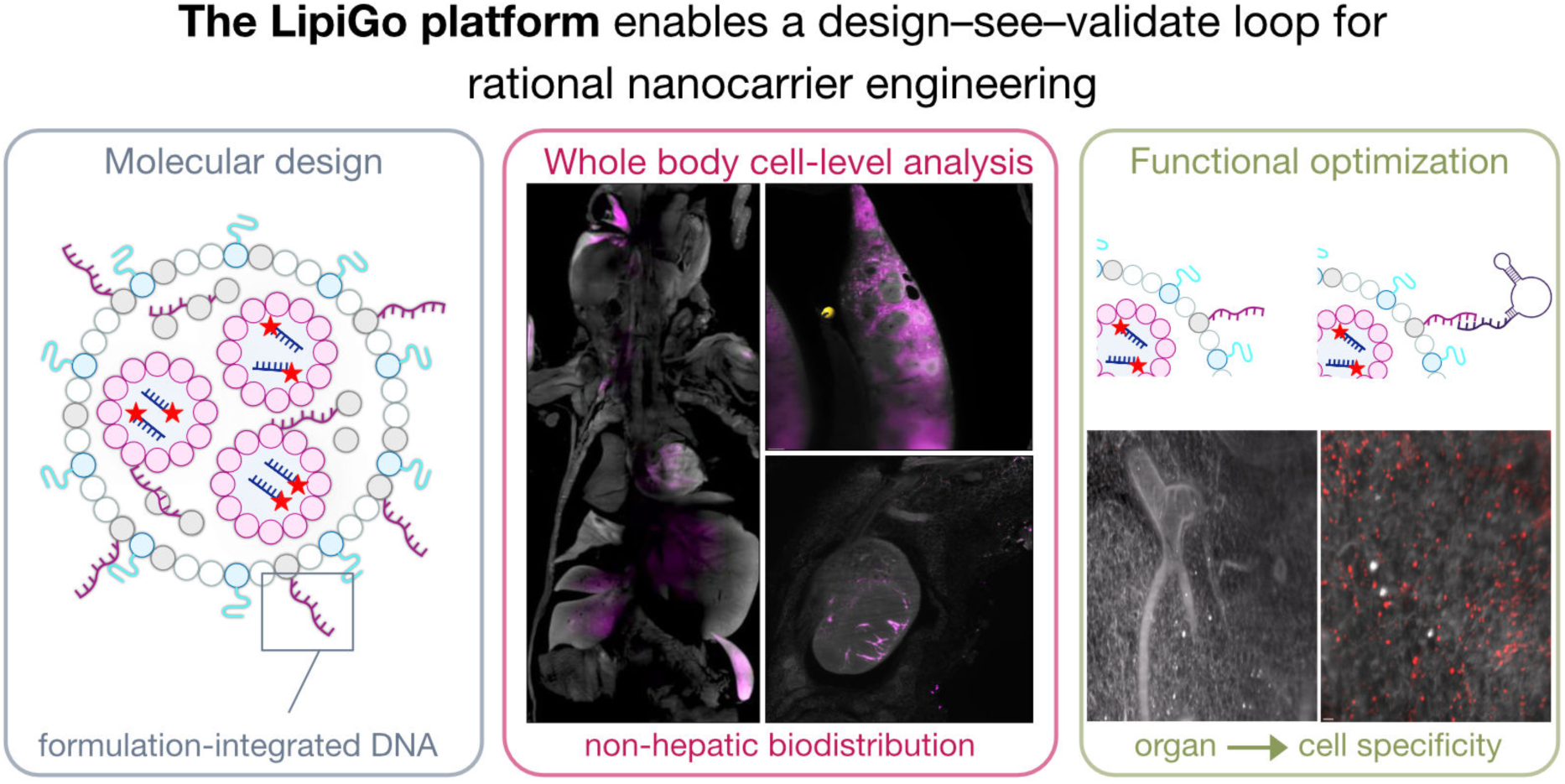

## 1. Introduction

Nucleic acid therapeutics offer opportunities for both prophylactic and therapeutic applications with the potential to address targets previously considered undruggable by conventional methods. Their broad clinical translation, however, remains constrained by targeted delivery.

Lipid nanoparticles (LNPs) represent one of the most clinically advanced vehicles for the delivery of nucleic acids as they combine high loading capacity, stability in circulation, and low toxicity. Their impact has become evident in approved medicines, including SARS-CoV-2 vaccines and a therapy for hereditary transthyretin-mediated amyloidosis, with numerous additional candidates currently in clinical trials.

Despite their advantages, intravenously administered LNPs intrinsically accumulate in the liver, which limits their utility in extrahepatic tissues and increases the risk of liver-related toxicity^1,2^. Although alternative administration routes can shift biodistribution toward different organs, precise delivery beyond the liver requires rational strategies to control nanoparticle tropism, which remains one of the central challenges in the field.

Efforts to achieve organ-, tissue-, and cell type-specific delivery with LNPs can be broadly categorized into passive and active strategies^3^. Varying lipid ratios, introducing lipids with distinct chemistries or expanding the number of constituent lipids reshapes the biomolecular corona that forms around LNPs upon exposure to biofluids, and thereby passively modulates interactions with immune receptors, cells, and tissues^4^. Active strategies, in contrast, involve decorating the particle surface with targeting ligands such as antibodies or aptamers^5,6^. Ligands are most commonly attached to LNPs via PEG-lipids, which protrude from the LNP surface^7^. However, PEG-lipids are designed to rapidly shed upon administration to facilitate cellular internalization and endosomal escape^8^. As PEG shedding typically occurs within minutes of administration, together with the strong intrinsic hepatic tropism of LNPs, efficient engagement of targeting ligands with their cognate receptors is often limited^9^. Thus, cell type-specific targeting following systemic delivery with active strategies has not been achieved *in vivo*^10^. Achieving sub-organ level specificity beyond the liver, therefore, requires overcoming dominant hepatic accumulation while simultaneously leveraging active targeting via surface-displayed ligands, highlighting the need for a vehicle design that integrates both passive and active targeting mechanisms.

Here, we introduce LipiGo, a lipid-based hybrid nanoparticles generated by partially substituting cholesterol with cholesterol-modified single-stranded DNA (ssDNA) that protrude from the particle surface. In this base formulation, the DNA strands alter particle physicochemical properties and redirect functional mRNA delivery from the liver toward immune organs. Beyond this intrinsic retargeting, the protruding DNA strands also serve as modular handles for ligand hybridization, thereby combining passive and active targeting in a single design and enabling cell-specific delivery, as demonstrated with a white adipocyte–targeting DNA aptamer.

## 2. Results and Discussion

### Design and characterization of LipiGo nanoparticles

To generate LipiGo, we partially substituted the cholesterol component in standard lipid nanoparticle formulations with a 26-base single stranded DNA (ssDNA) modified with a cholesterol moiety at the 3’ end. This modification results in nanoparticles with ssDNA protruding from their surface (**Fig. 1a,b**).

**Fig. 1.**
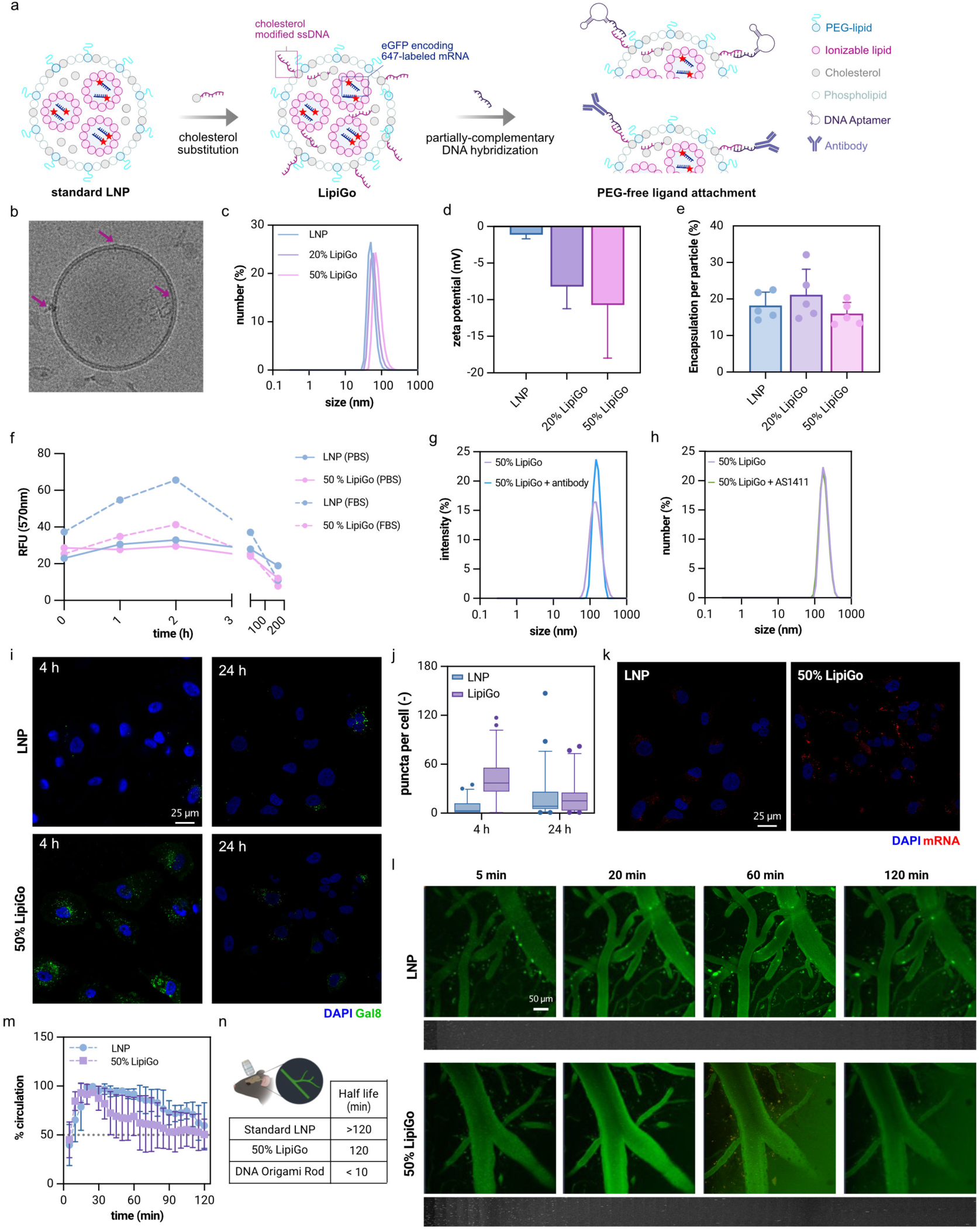
Design and characterization of LipiGo nanoparticles. **a)** Schematic of LipiGo design. Partial substitution of cholesterol with cholesterol-modified ssDNA enables ligand attachment via complementary base pairing. **b)** Representative Cryo-TEM image of LipiGo particle showing protruding DNA strands (arrows). **c)** Hydrodynamic size distribution of standard LNPs and LipiGo variants measured with DLS. **d)** Zeta potential measurements showing increased negative sharge with higher DNA substitution (N=5). **e)** mRNA encapsulation efficiency of LNPs and LipiGo variants (N=5). **f)** Stability of LNPs and LipiGo particles measured in PBS and serum supplemented media measured with FRET. **g,h)** Hydrodynamic size distribution of LipiGo before and after hybridization with IgG antibody (g) or AS1411 DNA aptamer (h). **i)** Representative confocal images of Gal8 recruitment assay in HeLa cells shows enhanced endosomal rupture at 4 h with LipiGo compared to LNPs. **j)** Quantification of Gal9 puncta per cell (N = 50 cell per condition**). k)** Representative confocal images of HeLa cells after 24 h incubation with LNP or LipiGo particles**. l)** Intravital two-photon microscopy images of LNPs and LipiGo in mouse vasculature at indicated time points. **m)** Quantification of circulation profiles showing slightly shorter half-life of LipiGo compared to standard LNPs. **n)** Summary of measured circulation half-lives comparing standard LNPs, LipiGo particles, and DNA origami particles^15^.

While generating the base LipiGo formulation, lipid ratios were adapted from Cheng *et al*.^11^ for efficient mRNA encapsulation and release. Particles were assembled by mixing an aqueous phase containing EGFP-encoding, fluorescently labeled mRNA with an ethanol phase containing DLin-MC3-DMA (23.8 mol %), PEG2000-C-DMG (4.3 mol %), DSPC (23.8 mol %), DMG-PEG(2000) (0.5 mol %), and cholesterol (47.6 mol %), with partial substitution of cholesterol by cholesterol-modified ssDNA at varying degrees.

Compared to conventional LNPs, LipiGo displayed altered physicochemical properties. Their hydrodynamic size increased slightly from ∼50 nm to ∼68 nm (**Fig. 1c**). Increasing the degree of cholesterol-ssDNA substitution from 20 to 50 % shifted the zeta potential towards more negative values (−8.2 ± 3 mV and −10.7 ± 7 for 20 % and 50 % LipiGo, respectively) (**Fig. 1d**). A similar change in zeta potential was observed across different DNA sequences and ionizable lipid chemistries (*i.e.*, ALC-1315 and SM1502), indicating that the effect is sequence and lipid chemistry independent (**Fig. S1a**). Across formulations, the polydispersity index (PDI) remained below 0.2 for all variants, indicating narrow and monomodal size distributions and RNA encapsulation efficiency was not significantly affected by addition of the DNA-cholesterol (**Fig. 1e**). Importantly, particles remain stable for at least 24 h in serum-supplemented media (**Fig. 1f**). Together, these results indicate that while preserving uniformity, stability and loading capacity, cholesterol-ssDNA protrusions provide a tunable parameter on nanoparticle features to alter physicochemical characteristics, *i.e.*, size and zeta potential, which are known to influence the interactions with biological material, affecting their biodistribution^4^.

Beyond tuning physical characteristics, the surface-exposed DNA handles of LipiGo provide modular sites for ligand attachment through complementary base pairing. To demonstrate this versatility, we conjugated two structurally distinct ligands, *i.e.*, an immunoglobulin G (IgG) antibody and a short DNA aptamer AS1411. The IgG antibody was first conjugated to a complementary DNA strand *via* maleimide chemistry, followed by hybridization to the LipiGo handle (**Fig. 1a**). The aptamer was extended with a short region complementary to the LipiGo handle strand and therefore could be directly hybridized to the handle DNA. Both ligand-conjugated LipiGo variants retained comparable physicochemical properties within an acceptable size range (**Fig. 1g,h**). Successful aptamer hybridization was confirmed by an increase in double-stranded DNA content together with a decrease in zeta potential (**Fig. S1c**). Importantly, this DNA-mediated conjugation strategy not only ensures controlled orientation and display of targeting ligands but also avoids the altered biodistribution and enhanced toxicity associated with complement activation, a phenomenon recently reported for nanoparticles functionalized with targeting antibodies using conventional chemistries such as dibenzocyclooctyne-mediated coupling^12^.

We next assessed the cellular compatibility and intracellular behavior of LipiGo *in vitro*. LipiGo particles did not induce detectable cytotoxicity within the tested concentration range (1-10 µg mRNA per mouse) as measured by cell viability assay in HEK cells (**Fig. S2a**). To evaluate endosomal escape and cytoplasmic delivery, we employed the Gal8 recruitment assay in HeLa cells, which are stably expressing Galectin8-mRuby3 fusion protein^13^. Upon endosomal membrane rupture, Gal8-mRuby3 binds to the inner lumen of the disrupted endosomes resulting in punctate fluorescent spots, indicating cytosolic release of the particle remnants and their cargo. We observed a significantly increased endosomal rupture at 4 h after LipiGo treatment compared to the standard LNPs (**Fig. 1i,j**). By 24 h, Gal-8 positive puncta in LipiGo treated cells decreased to levels comparable with those observed with standard LNPs, suggesting that LipiGo promotes more rapid endosomal escape. Consistent with this, cells incubated with LipiGo showed stronger fluorescent mRNA signal at 24 h relative to standard LNPs (**Fig. 1k**), indicating enhanced cytoplasmic retention and reduced intracellular degradation. These results suggest that LipiGo interacts differently with endosomal membranes or trafficking pathways, leading to more efficient release of its cargo. Since limited endosomal escape remains a key bottleneck in mRNA delivery, the accelerated endosomal rupture and sustained cytoplasmic mRNA signal with LipiGo highlights its potential to overcome this challenge.

Following in vitro evaluations, we next evaluated the half-life of LipiGo particles in bloodstream using in vivo two-photon microscopy. LipiGo exhibited a circulation half-life of ∼120 minutes, which was slightly shorter than that of the parent LNPs, suggesting faster tissue uptake. This finding is consistent with the enhanced cellular uptake observed *in vitro*, and fall falls within the range reported for other LNP formulations assessed with alternative techniques^14^ (**Fig. 1l-n**).

### LipiGo exhibits enhanced biodistribution to lymphatic tissues

To evaluate the *in vivo* biodistribution of LipiGo nanoparticles, we intravenously injected 1 µg mRNA/mouse (0.067 mg/kg mRNA) and analyzed particle distribution after 1 h using our in-house developed “Single Cell Precision Nanocarrier Identification (SCP Nano)” pipeline^15^. SCP Nano integrates whole-body tissue clearing of LNP/LipiGo administered mice, light sheet fluorescence microscopy (LSFM) imaging, and deep learning-based signal quantification at single cell-resolution for biodistribution mapping (**Fig. 2a**). As expected, intravenous administration of standard LNPs led to predominant hepatic accumulation, limiting their ability to effectively target other organs^2^. Using high resolution whole-body imaging, we observed LipiGo and LNP signals in other major organs including the spleen, lungs, and heart (**Fig. 2b**, **Fig. S3**). Some of these organ-specific signals would be difficult to capture using conventional techniques such as bioluminescence imaging (**Fig. S4b**).

**Fig. 2.**
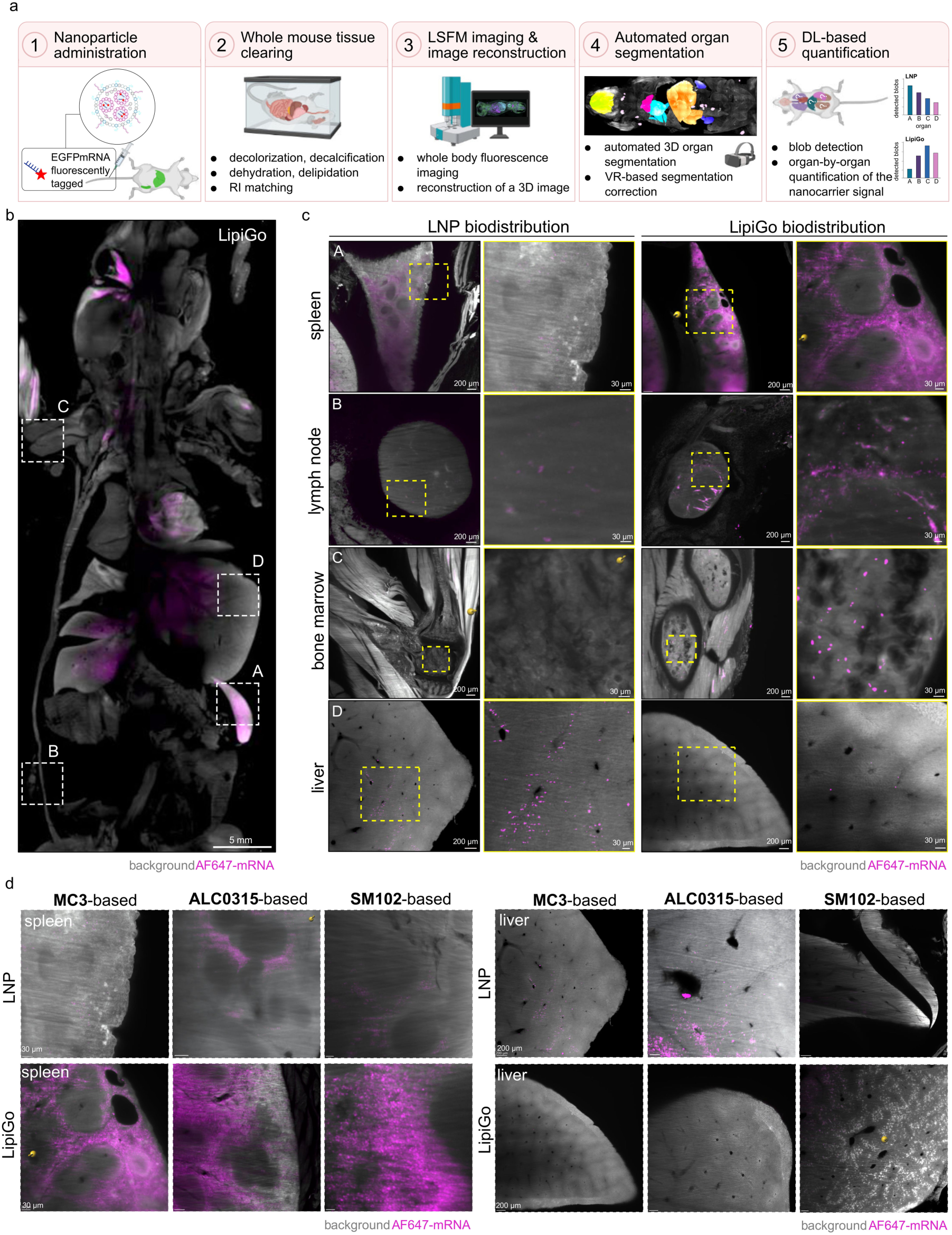
Whole-body mRNA biodistribution assessment of LipiGo particles assessed by SCP-Nano. **a)** Schematic of the SCP-nano workflow: (1) fluorescently labelled mRNA-loaded nanoparticle administration, (2) whole body tissue clearing, (3) light-sheet fluorescence microscopy (LSFM) imaging and 3D reconstruction, (4) automated organ segmentation with manual refinement, and (5) deep learning-based signal quantification. **b)** Whole-body 3D projection (600 µm slice) of a mouse 1 h after i.v. injection of LipiGo. **c)** Representative LSFM images of spleen, lymph node, bone marrow and liver, comparing biodistribution of standard LNPs and LipiGo formulations. Yellow dashed boxes indicate regions of interest (ROI) with close-up views, with particularly enhanced accumulation in immune organs for LipiGo. **d)** Comparison of spleen and liver biodistribution of nanoparticles formed with different base ionizable lipids (MC3, ALC0315, and SM-102), showing consistent spleen targeting across LipiGo variants.

Interestingly, LipiGo particles exhibited a pronounced tropism for lymphoid and immune associated organs, particularly in the spleen and lymph nodes, which are targeted less by conventional LNPs following intravenous administration (**Fig. 2b,c**). Standard LNPs and LipiGo particles also show differences in their intra-organ distribution (**Fig. 2c**). For example, liver distribution patterns for LipiGo particles were found to be more homogeneous and dispersed than for standard LNPs, which showed perivascular clustering. These differences likely reflect deeper tissue penetration associated with faster blood clearance as indicated by the lower signal at 60 minutes in the blood that was measured with intravitreal microscopy (**Fig. 1l-n**).

To test whether the observed shift in biodistribution was independent on the specific LNP formulation, we generated LipiGo particles using other clinically validated ionizable lipids, i.e., ALC-0315 (used in BioNTech’s COVID-19 vaccine) and SM-102 (used in Moderna’s Covid 19 vaccine). Physicochemical properties were similar across all LipiGo variants (**Fig. S1a**). Notably, incorporation of the DNA-lipid handle into the base formulation consistently led to a more uniform biodistribution pattern in the liver and to a substantial shift in tropism towards immune organs (**Fig. 2d, Fig. S3a**), supporting the compositional flexibility of the LipiGo platform.

We next evaluated whether sequence-level engineering of the DNA handle further modulates the biodistribution. We designed a DNA handle variant containing a CpG motif known to activate immune cells via Toll-like receptor (TLR9) ^16–18^. While the original LipiGo sequence was designed to be biologically inert, both sequences were identical in length and total GC content to control for structural effects. LipiGo particles formulated with the CpG-containing sequence (LipiGo-CpG), exhibited similar hydrodynamic size and zeta potential, ruling out colloidal effects on biological behavior (**Fig. S1b**). Following the SCP Nano pipeline, we found that LipiGo-CpG showed even stronger spleen tropism compared to the LipiGo with non-functional DNA handles, (**Fig. S3e**). These findings demonstrate that oligonucleotide sequence alone can be leveraged to tune targeting. Together, the findings demonstrate the modularity of LipiGo across different ionizable lipid backbones in maintaining immune organ targeting, as well as its tunability through DNA handle sequence level to further refine biodistribution, highlighting its flexibility for organ tropism.

### Quantitative organ and cell type profiling reveals LipiGo-mediated immune tropism despite residual hepatic distribution

Having qualitatively assessed particle distribution, we next performed deep-learning based quantification of organ-level targeting using automated segmentation of organs and mRNA signal (**Fig. 3a**). Based on whole-body images, we anticipated the major difference of LipiGo particles to be their reduced hepatic accumulation and increased localization to lymphatic organs (**Fig. 2a,b**). However, unbiased organ-by-organ quantification revealed interesting, yet, unexpected, findings. Even though the preferential accumulation of LipiGo in the spleen was validated, the visually perceived reduction in liver targeting was not supported quantitatively (**Fig. 3b**). Compared to LNPs, LipiGo particles exhibited slightly reduced lung accumulation, consistent with previous reports showing that LNPs formulated with cationic lipids preferentially localize to the lung whereas the more negatively charged formulations demonstrate spleen tropism^4^. Across all formulations, minimal particle accumulation was quantified in the brain, heart, and kidneys, and no significant differences were detected among LipiGo variants with varying degrees of DNA handle modification (**Fig. 3b**).

**Fig. 3.**
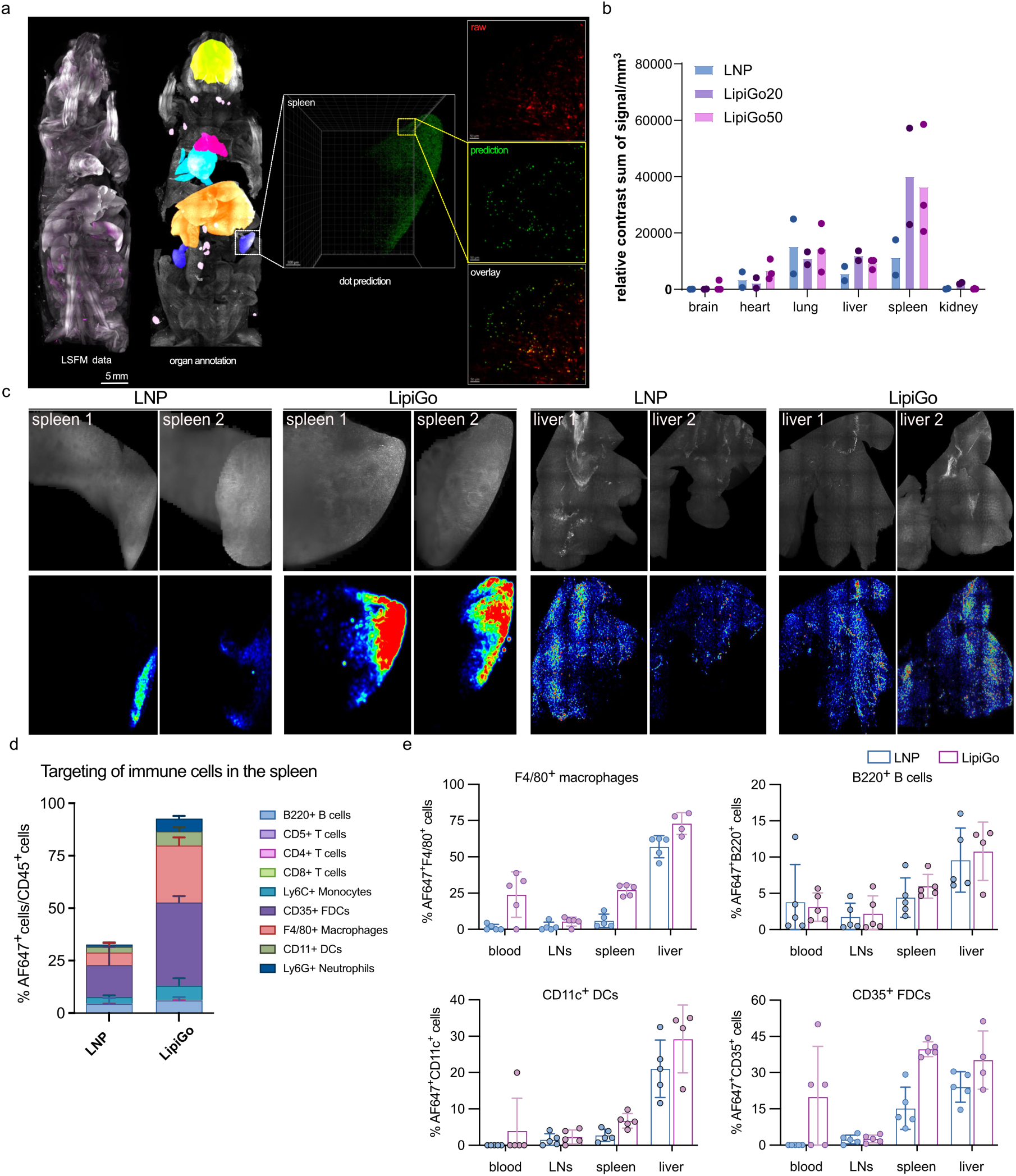
Quantitative profiling of LipiGo biodistribution at the organ and immune cell level. **a)** Light sheet fluorescence microscopy (LSFM)-based whole-body analysis workflow showing automated organ segmentation and nanoparticle signal prediction. Representative spleen region from a LipiGo-injected mouse is shown as raw (red), predicted (green), and overlay images. **b)** Quantification of relative nanoparticle accumulation across major organs 1 h post administration using SCP-nano analysis (N = 3). **c)** Representative SCP-nano quantified particle distribution density heatmaps for spleen and liver (N=2). **d)** Flow cytometry analysis of immune cells subtypes in the spleen (CD45^+^ cells) showing the percentage of LNP (mRNA-AF647^+^) positive cells in the respective immune cell populations (N=5). **e)** Quantification of AF647^+^ cells among F4/80^+^ macrophages, B220^+^ B cells, CD11c^+^ dendritic cells (DCs) and CD35^+^ follicular dendritic cells (FDCs) in blood, lymph nodes (LNs), spleen, and liver (N=5). Data are shown as mean ± SD.

Revisiting the raw images for a closer inspection provided insight into these observations. In LipiGo-injected mice, the mRNA signal in the livers appeared more sparsely distributed, whereas in standard LNP-injected mice, the signal was concentrated in focal regions as distinct patches near blood vessels. This perivascular clustering likely contributed to the apparent overestimation of liver enrichment by visual inspection alone, underscoring the importance of unbiased quantification. The more dispersed distribution pattern observed with LipiGo may reflect differences in particle kinetics. As shown in the circulation half-life experiments (**Fig. 1l-n**), fewer LipiGo particles remained in bloodstream at 60 minutes, suggesting more rapid tissue uptake. It is possible that, by the time of sacrifice, *i.e.*, 60 minutes post injection, LipiGo particles potentially penetrated deeper into liver parenchyma, leading to a more homogeneous and less visually prominent signal compared to the vascular-concentrated standard LNPs. This integrated view, together with the image-based quantification and kinetic data highlights the value of multimodal analysis in revealing subtle but biologically meaningful differences in nanocarrier behavior.

In addition, we generated liver and spleen biodistribution density maps to facilitate direct comparison of accumulation patterns between standard LNP and LipiGo formulations in terms of liver to spleen tropism. These density heat maps further illustrated these differences, with LipiGo-injected animals showing fewer high-density patches in the liver and stronger accumulation in the spleen (**Fig. 3c**). The enhanced spleen targeting of LipiGo was consistently validated by SCP-nano, as demonstrated both in the representative heat maps (**Fig. 3c**) and in corresponding quantitative dot counts at the organ level (**Fig. 3b**).

With LipiGo showing a strong tropism for lymphoid organs, we next investigated the immune cell types targeted across key immune-relevant tissues including blood, lymph nodes, spleen, and liver. Flow cytometric profiling of 20 distinct immune cell subsets revealed a clear shift in cellular tropism compared to standard LNPs (**Fig. 3d, Fig. S5**). Notably, LipiGo demonstrated enhanced uptake across several immune cell types, particularly in secondary lymphoid organs. In the spleen, LipiGo increased the percentage of mRNA homing immune cells from 26.51% to 65.16% with the most prominent contributions from antigen presenting cells such as CD35^+^ follicular dendritic cells (5.8-fold), F4/80^+^ macrophages (11.3-fold), and CD11c^+^ dendritic cells (2.4-fold) (**Fig. 3d,e**). T-cell delivery also improved with higher frequencies of both CD4+ and CD8+ subsets labeled in the spleen and lymph nodes, despite overall lower levels. High targeting rates of both the antigen presenting and effector cells at these critical sites suggests a potential utility of LipiGo particles to trigger and shape specific immune responses. Furthermore, beyond adaptive immunity, LipiGo broadened innate immune engagement with elevated signals observed in Ly6C⁺ monocytes and Ly6G⁺ neutrophils across peripheral organs. By contrast, standard LNPs showed substantially lower mRNA delivery to these immune subsets in the spleen, and remained largely confined to liver-resident cells. Notably, even within the liver, LipiGo outperformed standard LNPs for most immune cell subtypes, with strong increases in targeting efficiency in CD4^+^ T cells and Ly6C⁺ monocytes and Ly6G⁺ neutrophils. Together, these results highlight the potential of LipiGo to engage both innate and adaptive immune cells within immunologically active tissues.

### Functional delivery of mRNA to the immune cells with reduced hepatic expression

Having demonstrated through whole body mRNA cargo imaging that LipiGo enhances tropism to lymphatic organs, we next investigated whether these particles enable functional delivery, *i.e.*, successful cytoplasmic mRNA release and translation to protein. Mice were injected intravenously with LipiGo particles carrying EGFP mRNA (0.067 mg/kg) and sacrificed 48 h later. EGFP expression was detected with nanobody staining using the nanoDISCO whole body clearing method ^15^.

In contrast to the biodistribution data, functionality readout revealed significantly reduced EGFP expression in LipiGo-administered mouse livers (**Fig. 4a,b, Fig. S4a**). Moreover, we observed different protein expression distribution pattern within the liver compared to standard LNPs. While LNPs induced concentrated, perivascular clusters of EGFP, LipiGo-treated livers exhibited minimal and more diffuse signal (**Fig. 4b**). By contrast, LipiGo still induced robust EGFP expression in lymphoid tissues, with spleen expression markedly surpassing that of LNPs. In addition to the spleen, higher EGFP-signals are also seen in thymus and in the lymph nodes when the mRNA is delivered with LipiGo rather than with standard LNPs. Thus, although LipiGo distributes systemically, efficient mRNA translation is selectively enriched in immune organs.

**Fig. 4.**
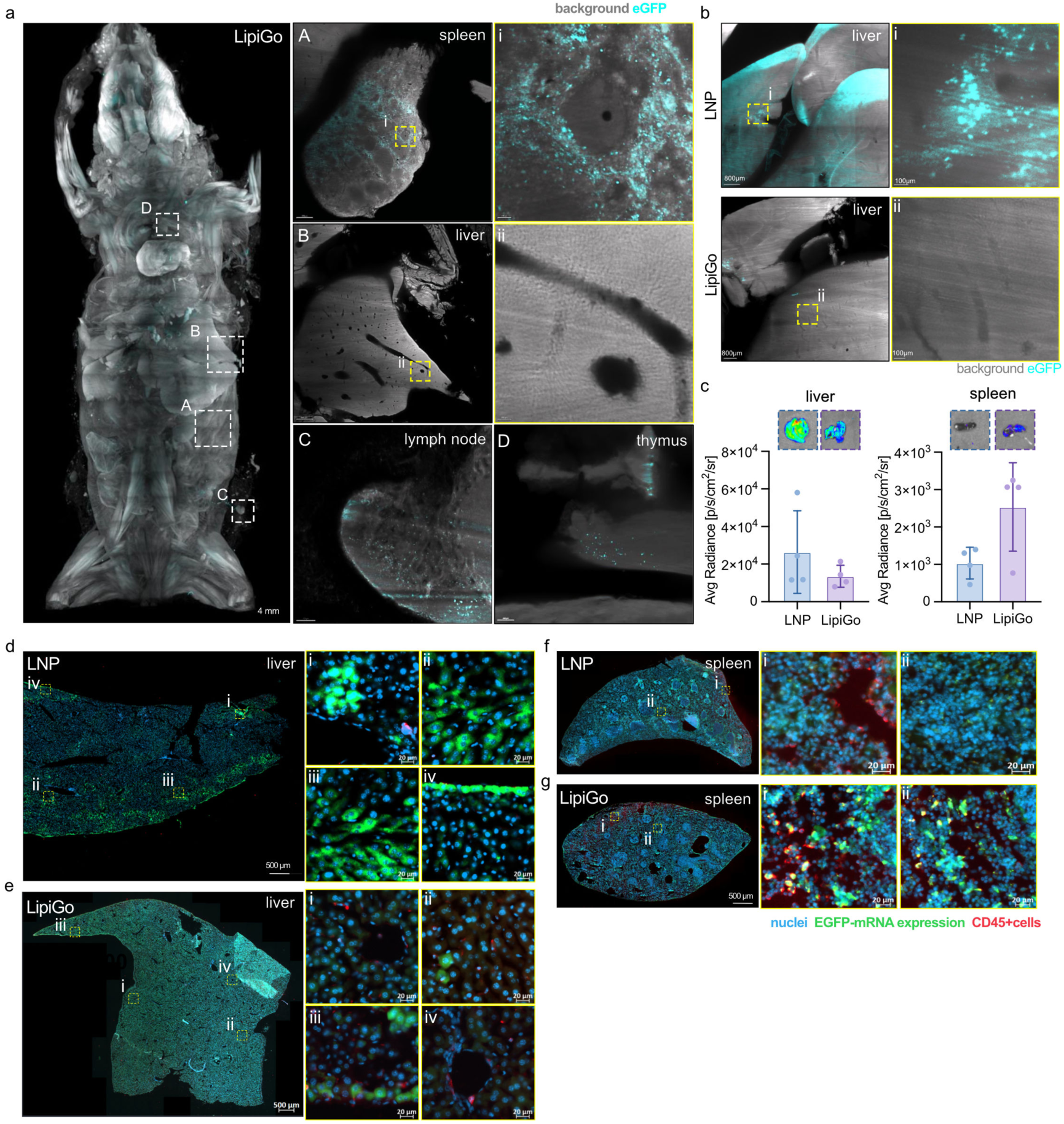
LipiGo enables functional mRNA delivery to immune cell rich organs. **a)** Whole-body 3D projection (600 µm) of a mouse 48 h post i.v. injection of LipiGo delivering EGFP mRNA. Dashed boxes indicate ROIs in the spleen, liver, lymph node and thymus. **b)** EGFP expression in livers of mice treated with standard LNPs or LipiGo, demonstrating reduced hepatic expression with LipiGo. **c)** Bioluminescence imaging and quantification of FLuc mRNA expression in liver and spleen 24 h post-injection of standard LNP and LipiGo (N=3). **d-g)** Immunofluorescence staining of fresh frozen tissue sections of liver (d,e) and spleen (f,g) (10 µm) showing EGFP expression (green), DAPI nuclear staining (blue), and CD45^+^ immune cells (red).

To validate this organ tropism with an orthogonal readout, we performed bioluminescence imaging by loading nanoparticles with FLuc mRNA (0.067 mg/kg). Quantitative analysis of signal in the spleens and livers confirmed the enhanced expression in the spleen and confirmed the functional liver detargeting effect in LipiGo-treated animals (**Fig. 4c, Fig. S4b**).

To resolve the expression at the cellular level, we carried out immunofluorescence staining of fresh frozen tissue sections from the livers and spleens (**Fig. 4d-f**). Consistent with whole-body imaging, standard LNPs induced strong EGFP expression in the liver, whereas LipiGo-treated livers showed minimal signal. In the spleen, however, LipiGo exhibited robust and broadly distributed protein expression (**Fig. 4f**). Notably, unlike the clustered expression patterns seen in standard LNP-treated livers, LipiGo-mediated expression appeared more diffuse and less concentrated. Immunostaining confirmed that LipiGo particles were internalized to greater extent by CD45^+^ immune cells as compared to standard LNPs, and resulted in slightly elevated number of cells received functional delivery in these populations (**Fig. S4c**). These findings again confirm the flow cytometry results, providing a direct functional readout of protein expression in immune cell subsets.

### LipiGo elicits minimal systemic response and forms a Angpt1-rich protein corona

Having established functional delivery of mRNA to immune tissues, we next evaluated the in vivo safety profile of LipiGo by assessing systemic cytokine responses and serum protein composition. Cytokine levels were profiled 48 h post-injection (0.168 mg mRNA/kg mouse) using a 13-plex flow-cytometry based immunoassay.

Except for IL-1α, cytokine concretions remained below the technical limit of quantification for both standard LNPs and LipiGo, suggesting that no major systemic inflammatory response was triggered (**Fig. S2b**). Key inflammatory mediators commonly associated with acute nanocarrier toxicity, including IL-6, TNF-α, and IFN-γ were undetectable in both groups. Notably, the absence of IL-6 induction is particularly relevant given the presence of ssDNA on the LipiGo surface, which could potentially engage TRL9 (**Fig. S2b**).

Next, we measured markers for metabolic stress and liver toxicity in mouse serum after 1h of LNP/LipiGo circulation, injecting a high dose of 0.67 mg mRNA/kg mouse (**Fig. S2c**). LipiGo and LipiGo-CpG showed a modest but non-significant increase in glucose and lactate compared to standard LNPs and untreated controls, hinting to a general inflammatory response reaction after the injection of the particles. However, since the cytokine panel 48h after injection does not show adverse effects, the inflammatory response seems to be small and of short duration. All liver stress and damage markers (AST, ALT, KETONE) were similar to the negative control for all particles - indicating that neither of the particles causes hepatotoxicity (**Fig. S2c**). Together, these findings demonstrate that LipiGo maintains a favorable safety profile, with minimal systemic immune activation and no signs of acute toxicity, even at elevated doses.

Upon exposure to biological fluids, nanoparticles rapidly adsorb a layer of biomolecules primarily composed of proteins on their surface, forming a biomolecular corona that critically influences their colloidal stability, biodistribution, and organ tropism^19–22^. To determine whether an altered corona composition contributes to the distinct tropism of LipiGo particles, we performed comparative proteomic profiling of the protein coronas formed on LipiGo and standard LNPs using LC-MS/MS (**Fig. S6a**). Principal component analysis (PCA) revealed clear separation between the protein corona profiles of LipiGo and standard LNPs, independent of CpG-rich motif inclusion, suggesting that the DNA-handle modification consistently alters protein adsorption patterns (**Fig. S6b**).

Interestingly, among the differentially enriched proteins, Angiopoietin-1 (Angpt1) was consistently and significantly enriched in all LipiGo coronas independent of the ionizable lipid used in formulation (**Fig. S6c**). The consistent enrichment of Angpt1 in the LipiGo protein corona may contribute to enhanced immune organ targeting by promoting interactions with Tie2-expressing endothelial cells abundant in lymphatic tissues, including lymph nodes and spleen^23^. These interactions could facilitate nanoparticle retention or uptake within these organs. Moreover, Angpt1 is known to regulate endothelial cell stability and vascular permeability. Its significant enrichment on the nanoparticle surface could modulate endothelial barrier properties, potentially enabling more efficient nanoparticle extravasation from the bloodstream into tissues and explaining the faster blood clearance and more tissue distribution of LipiGo that we observed. These combined effects, *i.e.*, enhanced receptor mediated interactions and altered vascular permeability may contribute to the observed redistribution of LipiGo nanoparticles toward lymphatic organs and the faster and deeper tissue penetration.

### Aptamer functionalization of LipiGo enables cell specific targeting *in vitro* and *in vivo*

So far we showed that the integration of ssDNA into an LNP formulation passively altered their *in vivo* fate. However, beyond changing the physicochemical characteristics, these DNA strands also provide a versatile platform for active ligand attachment *via* DNA hybridization, enabling straightforward conjugation of diverse targeting ligands including antibodies or their fragments, peptides, and DNA/RNA aptamers.

To demonstrate the utility of DNA handles for active targeting, we functionalized LipiGo particles with DNA-based aptamers by hybridization to complementary DNA handles pre-incorporated in the formulation (**Fig 1a**). Upon aptamer hybridization, we observed a remarkable shift towards a more negative zeta potential (**Fig. S1c**). Although minor non-specific adsorption was detected on control LNPs lacking the DNA handle, the pronounced changes in these physicochemical properties confirmed specific and efficient aptamer conjugation.

First, we evaluated the targeting performance of aptamer-functionalized LipiGo *in vitro* using differentiated white and brown adipocytes. For these experiments, we selected a novel DNA aptamer AS1 specifically targeting the Asc-1 protein^24^, which is highly expressed on white adipocytes but not on brown adipocytes.

Confocal microscopy revealed substantial internalization of AS1-LipiGo particles carrying fluorescently labeled mRNA into the cytoplasm of white adipocytes in an incubation period of 40 minutes (**Fig. 5a**). Cargo mRNA was localized as clusters within intracellular compartments, suggesting endosomal trafficking. In contrast, untargeted LipiGo particles and control LNPs incubated with free AS1 aptamers exhibited only weak, non-specific surface association with negligible cellular uptake (**Fig. 5b**, upper panel). To confirm cell type specificity, the same formulations were incubated with brown adipocytes lacking the target surface protein. As expected, all formulations including the targeting AS1-LipiGo, showed minimal cellular uptake (**Fig. 5b**, lower panel). These results collectively demonstrate that active functionalization of LipiGo particles enables the specific internalization of mRNA into target cells. Having confirmed specific uptake *in vitro*, we next evaluated whether the active targeting strategy translates to tissue-specific delivery *in vivo*. Whole body clearing and imaging revealed very strong accumulation of AS1-LipiGo in the white adipocyte fat tissue (WAT) close to the injection site (**Fig. 5f**). Next, to assessed the functional delivery. AS1-LipiGo particles encapsulating luciferase-encoding mRNA were administered subcutaneously into mice. Live animal bioluminescence imaging (**Fig. S7a**) and subsequent *ex vivo* analysis of harvested organs 24 h post-injection revealed highly localized and robust luciferase expression predominantly within the white adipose tissue proximal to the injection site (inguinal WAT) (**Fig. S7b**). Importantly, this expression was significantly higher compared to that from untargeted LipiGo controls, indicating the tissue target specificity of AS1-LipiGo particles. While minimal off-target expression was detected in the liver, spleen, and kidneys, the expression in distal WAT depots (*e.g.*, interscapular) was also negligible (**Fig. 5g**). This biodistribution profile suggests that either most of the particles were readily taken up by the inguinal WAT already before reaching other parts of the body, or that biodistribution throughout the body takes longer than 24 h when administered subcutaneously. In conclusion, these findings demonstrate that the LipiGo platform enables the modular attachment of functional aptamers for cell-specific delivery and expression of mRNA cargo *in vitro* and *in vivo*, establishing a promising and versatile approach for targeted nanomedicine.

**Fig 5:**
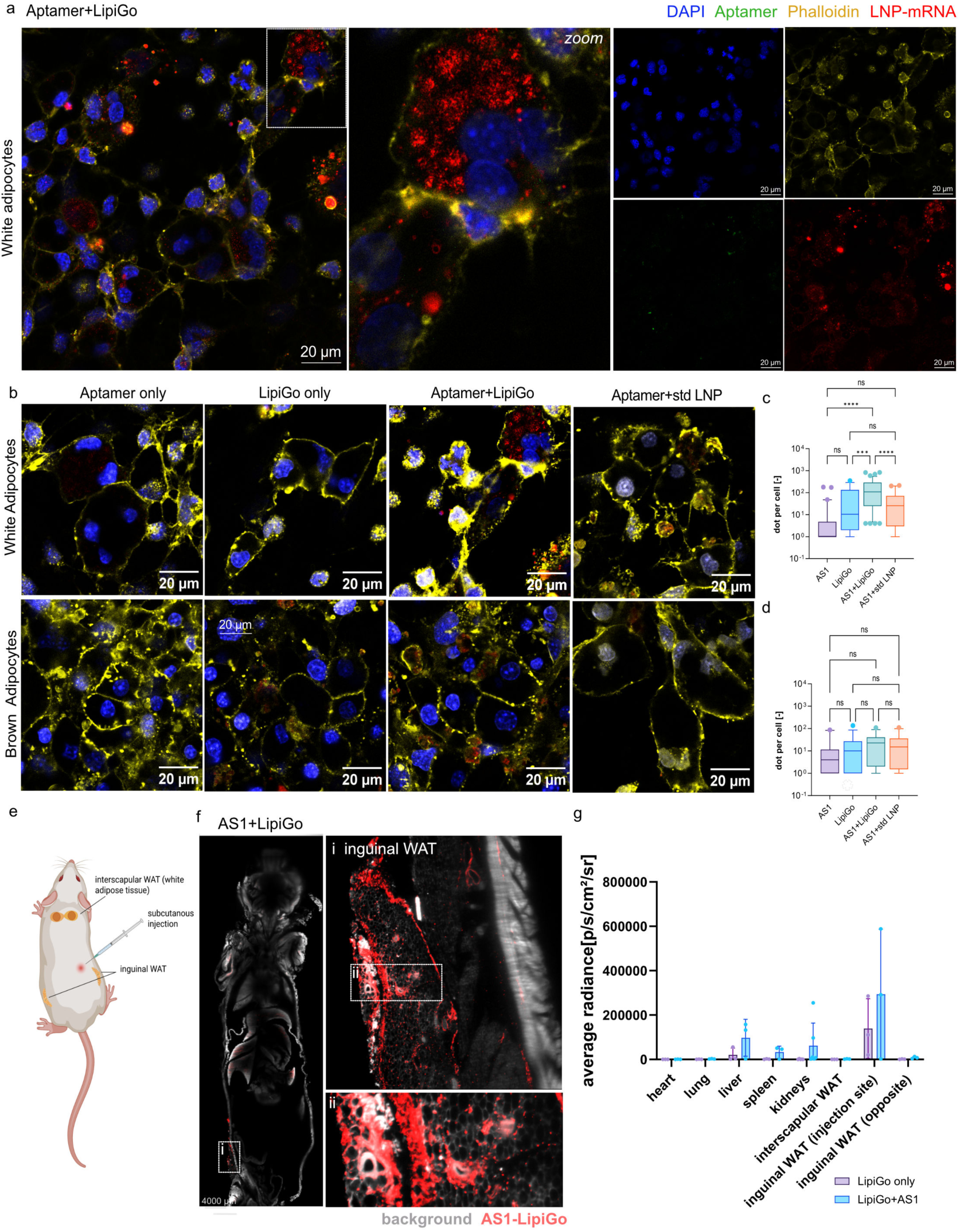
Specific targeting and functional mRNA delivery to white adipocytes *in vitro* and *in vivo*. **a)** Representative confocal microscopy image of differentiated white adipocytes after incubation with LipiGo particles conjugated with the Asc1-surface protein-targeting aptamer (AS1-LipiGo). The particles encapsulate labeled mRNA (red) and are functionalized with a FAM-labeled aptamer (green). The image shows significant internalization and co-localization of both components within intracellular compartments. Cell nuclei are stained with DAPI (blue). **b.** *In vitro* specificity assessment. The top row shows white adipocytes incubated with control formulations: non-conjugated AS1 aptamer, untargeted LipiGo, and standard LNPs lacking the DNA handle but unspecifically mixed with AS1. Only the fully conjugated AS1-LipiGo particles show significant uptake. The bottom row shows the same set of experiments performed on brown adipocytes, which lack the insulin receptor. **c-d.** Quantification of the mRNA signal inside the white (c) and brown (d) adipocytes. One-way ANOVA with Tukey’s multiple comparison test with a single pooled variance is employed. **e.** Schematic overview of the *in vivo* study design. Mice were subcutaneously (s.c.) injected with either AS1-LipiGo or untargeted LipiGo particles, to evaluate targeted delivery and functional protein expression. **f.** Whole body imaging of a tissue cleared mouse showing the biodistribution of AS1-LipiGo after 1 h circulation (left) with a zoom in to inguinal WAT (right). **g.** *Ex vivo* quantification of functional mRNA delivery. Bioluminescence signal from luciferase expression was measured in harvested organs 24 hours post-injection. The quantification reveals that AS1-LipiGo particles (**d**) mediate significantly higher luciferase expression in the target inguinal white adipose tissue (WAT) compared to the untargeted LipiGo control particles (**e**), with minimal off-target expression observed in other organs. Data are presented as mean ± s.d. (n=3 mice per group).

## Conclusions

The ability to direct nucleic acid nanocarriers to specific organs, tissues and cell types is a central step towards expanding their therapeutic utility. In this study, we demonstrate LipiGo as a flexible platform that combines passive and active ligand-mediated targeting within a single nanocarrier design, By embedding DNA handles into the LNP framework, LipiGo combines altered characteristics driving redistribution from the liver to lymphoid organs with modular functionalization through DNA hybridization. This dual strategy expands the therapeutic reach of LNPs well beyond hepatic applications. LipiGo enabled selective and functional mRNA delivery to immune cell-rich organs while reducing hepatic expression, and further demonstrated utility as active targeting nanocarrier, achieving receptor specific delivery to white adipocytes *in vitro* and *in vivo*. The ability to unite systemic detargeting, lymphoid tropism and customizable ligand attachment highlights LipiGo as a programmable nanocarrier platform. This modular strategy could enable rapid customization of nanocarriers across diverse therapeutic areas - providing a blueprint for programmable drug delivery systems. Future studies will be directed at translating LipiGo-based constructs into therapeutic applications.

## Methods

### LNP/LipiGo preparation

LNPs/LipiGo particles were prepared using the ethanol dilution method. All lipids with specified molar ratios (Dlin-MC3-DMA/DSPC/cholesterol/PEG-DMG at 47.6:23.8:23.8:4.8 %) were dissolved in ethanol absolute (<99.8%), and mRNA was dissolved in 50 mM acetate buffer (pH 4.0). Those two solutions were mixed at an aqueous-to-ethanol volume ratio of 3:1 to a final N:P ratio of 6 and then incubated for 10 min at room temperature. The final LNP formulations were dialyzed (MWCO 3.5 kDa) against sterile filtered PBS for 1 h per 100 ul LNP volume to remove the ethanol and neutralize the pH. Standard LNPs were formulated using the molar ratios above. For LipiGo particles, a certain molar percentage (10%, 20% or 50%) of the cholesterol was subsidized with cholesterol bound ssDNA (26nt long) and included in the lipid phase before formulation, the particles were then dialyzed with dialysis tubes with a MWCO of 12kDa.

### Physicochemical characterizations of LNP/LipiGo particles

#### DLS and Zeta potential measurements

The hydrodynamic diameter, polydispersity index (PDI) and zeta potential of the LNPs were measured by DLS (Malvern Zetasizer Pro). For measuring the size, the particle solution was diluted 1:50 with sterile filtered PBS and for measuring the zeta potential particles were diluted with nuclease free water. Plots were generated with GraphPad Prism and show the mean size (nm) of triplicates ± SD.

#### Attachment of targeting ligands to LipiGo particles

Attachment of aptamers: Aptamers with a bridge complementary to the ssDNA protruding from the LipiGo particles was folded by heating at 95°C for 5 min and then cooling down on ice until further use. LipiGo particles and aptamers were mixed at a molar ratio of 1:1 (binding sites on particles: aptamer) in PBS and then hybridized on a temperature ramp in a thermocycler at 37°C-20°C, −1°C per minute. Unbound aptamers (30 kDa) were removed using 50 kDa Amicon filters (Sigma-Aldich, UFC505008), that were pre-wetted with 0.1M PBS and then filled with the LipiGo-aptamer product and spun down at 8000 rcf for 10 minutes. 3 PBS wash steps followed, each for 5 min at 8000rcf, before inverting the filter into a new collection tube and spinning down the purified product

Attachment of protein-based ligands: Protein based ligands, such as antibodies, were attached to the thiolated complementary ssDNA strand of the LipiGo anchored handle. First, antibodies were functionalized with maleimide groups using SMCC chemistry as described in the reference^25^. Briefly, antibodies were dissolved in PBS (pH 7.2) and reacted with a 20-fold molar excess of Sulfo-SMCC (A39268, Thermo Fisher Scientific), dissolved in DMSO (final DMSO with antibody <10% v/v), for 30 min at room temperature. The activated antibody was purified using desalting columns (PD Spintrap-25, GE28-9180-04, Sigma-Aldrich) into PBS containing 1 mM EDTA. Separately, thiol-modified oligonucleotides were reduced by incubation with a 100-fold molar excess of TCEP (T2556, ThermoFisher Scientific) for 20 min at room temperature. Excess TCEP was removed by ethanol precipitation at −80 °C. For conjugation, reduced oligo-SH was added to maleimide-activated antibody at a 10:1 molar ratio (oligo:antibody), and the mixture was incubated overnight at 4 °C. Final conjugates were purified using 50 kDa molecular weight cut-off centrifugal filters (Amicon Ultra) to remove unbound oligo.

The antibody-oligonucleotide product was then hybridized to the LipiGo platform the same way as the aptamer as described above.

#### Single particle profile (SPP) analysis of particles containing mRNA

SPP measurements were carried out as described by Sych et al.^26^ To measure the number of particles carrying mRNA cargo, cargo and lipid shell had to be labeled in different fluorescent wavelengths. Particle cargo (mRNA) was labeled with AF-647 and the lipid particle core was labeled with Fast-DiO(D3898, Invitrogen). Single Particle Profiling was performed using the setup for confocal acquisition („spot scan”) at a Zeiss LSM 780 microscope. A 488 nm argon ion laser was used for FastDiO, whereas a 633 nm He−Ne laser was used for Alexa 647. A 40×1.2 NA water immersion objective was used to focus the light. The fluorescence emission was detected by GaAsP spectral detector in integration mode. The emission detection windows were set as 490 –560 for FastDiO and 650 –700 for AF-647. Intensity traces were recorded for 10 minutes, in total 5 measurements per sample. Curves were then analysed using the python program “Py Profiler”, from the reference above.

#### Serum stability

The stability of the particles in serum (from mice) was tested by measuring the intensity of the fluorescent resonance energy transfer (FRET) pair, DiO/DiI-with whom the particles were labeled. A 1:10 mixture of LNPs : mouse serum or LNP:PBS (control) was incubated at 37°C and samples taken at certain timepoints to be measured on a spectrometer (Varioskan). The fluorescence intensity reduction was measured at excitation and emission wavelengths of 480 nm, 530 nm (DiO), and 580 nm (DiI), respectively, to calculate the FRET efficiency. Plots were generated with GraphPad Prism, n=3, mean is shown.

### Cryo-TEM imaging of particles

The preparation of LNPs was done immediately before examination and stored in the fridge until the measurements. 4.5ul of a concentrated LNP formulation (lipid concentration 2.3 mg/mL) was placed on a thin carbon-reinforced copper grid for 6 min. The grid was blotted with a filter paper (blot time 4s, blot force 0) to remove the excess of liquid to form a thin film. The samples were vitrified rapidly in liquid ethane at −180 °C. Cryo-TEM measurements were performed using the Glacios TEM from ThermoFisher.

### Immunological profiling of injected LNPs/LipiGo particles

#### Plasma protein analysis

Mouse plasma of particle injected mice was collected after 1h or 48h of injection and stored in −80°C until further analysis with a Thermo Scientific Konelab analyzers to measure the concentration of plasma proteins.

#### FACS-based cytokine panel analysis

The LegendPlex immune panel (740929, BioLegend) allows for simultaneous quantification of 13 key targets essential for immune response such as IL-4, IL-2, CXCL10 (IP-10), IL-1β, TNF-α, CCL2 (MCP-1), IL-17A, IL-6, IL-10, IFN-γ, IL-12p70, CXCL8 (IL-8), and Free Active TGF-β1.6week old female BALB/c mice were intravenously injected with different formulations (PBS only, std LNP, LipiGo particles) containing 10 µg mRNA/mouse. After 1 h, the blood serum was obtained and used to measure the concentration of pro-inflammatory cytokines using the cytokine panel kit. Mouse Cytokine Panel 2 kit.

### Protein-corona analysis

Blood serum from wild-type Balb/c mice was collected retro-orbitally (in non-coated microcentrifuge tubes), the blood was left to clot at room temperature for 30 min and then spun down for 15 min at 2,000 r.p.m. at 4 °C to retrieve the serum (supernatant). LNPs (50 µl total volume each as described above, encapsulating 1 µg of unlabeled mRNA per formulation) were incubated with mouse serum for 15 min at 37 °C at a 1:1 volumetric ratio. A 0.7-M sucrose solution was prepared by dissolving solid sucrose in MilliQ water. The LNP/serum mixture was loaded onto a 300-µl sucrose cushion centrifuged at 15 300g and 4 °C for 1.5 h. The supernatant was removed, and the pellet was washed 3 times with 1× PBS, by spinning at 15 300g and 4 °C for 5 min and carefully removing the supernatant. The protein-covered LNPs were then kept dry at −20 °C upon mass spectrometry analysis.

To prepare the particles for MS-analysis, they were resuspended in 4% SDS-Tris-HCl buffer (4% SDS, 10 mM TCEP, pH 8.5) and incubated at 95 °C for 5 min, followed by sonication for 15 min at 4 °C. Subsequently, 2% SDC buffer (10 mM TCEP, 40 mM CAA) was added, and samples were heated again at 95 °C for 10 min. Proteins were precipitated by adding 3 µl of a 1:1 SpeedBeads mix and ethanol (final concentration 75%) under agitation for 5 min. Beads were immobilized using a magnetic rack, washed twice with 80% ethanol, and dried in a SpeedVac. Samples were digested overnight at 37 °C in 50 µl of 50 mM TEAB containing 0.4 µg Trypsin-LysC and 0.02% LMNG. Peptides were then cleaned using a modified Evotip PURE protocol in a 96-well format, at the end of which they were dried and resuspended in 0.1% TFA containing 0.015% DDM for MS analysis. Finally, peptides (50 ng) were analysed by LC–MS/MS on an EASY-nanoLC 1200 system (Thermo Fisher Scientific) coupled to a timsTOF SCP mass spectrometer (Bruker Daltonik) via a CaptiveSpray ion source. Data were acquired in DIA-PASEF mode with 100 ms ramp and accumulation times. For comparative analysis, 50 ng of peptide was injected onto a 5.5-cm μPAC Neo HPLC column (Thermo Fisher Scientific) and analysed using an 80-min gradient at a flow rate of 250 nl min⁻¹.

Data were analyzed using scanpy (v. 1.10.2) and anndata (v. 0.10.8) in Python 3.11. All proteins expressed in less than half of the samples in each group were filtered out, the data was log-transformed and normalized per sample. The missing values were input using KNNImputer (n_neighbors=5) from sklearn package (v. 1.5.1). With scanpy’s dendrogram function scipy’s hierarchical linkage clustering was calculated on a Pearson correlation matrix over groups which was calculated for 50 averaged principal components. Differential expression analysis was conducted using Scanpy’s method ‘rank_genes_groups’ with method set to ‘t-test’. A threshold of p < 0.05 and |log fold change| > 1.0 were applied to identify differentially expressed proteins (DEPs). These DEPs were subsequently visualized using volcano plots.

### Cellular uptake and endosomal escape experiments

HeLa or HeLa-Gal8-mRuby cells were seeded on µ-Slide 18-well glass bottom plates (81817, ibidi) with a density of 10000 or 7000 cells/well 1 day prior to particle exposure. LNP/LipiGo particles were added at an end concentration of 40 nM per well and cells were washed and fixed after 1h/4h or 24h.

For samples where lysosome co-staining was desired, AF488-mRNA containing particles were used. HeLa cells were stained with lysotracker DeepRed (L12492, ThermoFisher) at an end concentration of 50 nM. 5 min before ending the experiment, cells were stained with Hoechst 33342 (H3570, Invitrogen) at 5ug/ml in PBS 5 min before ending the experiment. Cells were gently washed 3x with warm DPBS to remove excess dyes and then imaged live at an SP5 Leica confocal microscope, image analysis was done with ImageJ.

To study uptake in recycling endosomes, Rab11 stainings were conducted in HeLa cells, who were subjected to std LNPs/LipiGo particles. After the incubation of particles with cells, cells were gently washed 3x with DPBS, fixed with 4% PFA-PBS at RT for 10 min and then permeabilized for 5 min with 0.1% Triton X-100. Blocking was done with 3% BSA (Carl Roth, 8076.2) for 30 min at RT, after which the cells were stained with 5ug/ml Rab11A-Alexa Fluor 488 (clone 3H18L5, ThermoFisher) (in blocking buffer) at RT for 1h. Dye removal was described as above, before fixing the cells using cold 4% PFA (1176205000, Morphisto) for 10 min at room temperature. Cells were finally washed again (first wash step including Hoechst) and then imaged as described above.

To visualize endosomal escape, AF647-mRNA containing particles were incubated with HeLa-mRuby cells, who emit fluorescence at 592 nm upon endosomal rupture. Cells were washed and fixed as described above and then imaged on a confocal microscope. Image representation and “puncta per cell” analysis was done with ImageJ.

### Aptamer-guided cell-culture targeting of white adipocytes

Standard LNPs and LipiGo(20) particles, decorated with targeting aptamers, were formulated as described above. Cells were seeded in 8-chamber slides (LabTek II) 9 days prior to the experiment. Differention was induced the next day (D0) with induction medium (DMEM, 10% FBS, 1% PenStrep, IBMX-KOH, Indometacin, Dexamethason, T3, Rosiglitazon and Insulin). At D2, medium was chagend to differentiation medium (DMEM, FBS, PenStrep, Rosi, T3, Insulin) and at D5 to differentiation medium without Rosi.

40 min before adding the particles, medium was changed to 130 uL fresh blocking medium ( dif. medium containing 100 mg/ L yeast tRNA). 50 ul of the particle solution was added to the wells, so that the resulting concentration of mRNA would be 40 nM in each well. For blank samples, 50 µL of sterile PBS was added to each well. After 40 minutes of particle incubation at 37°C, cells were washed with ice cold PBS before being fixed with 4% PFA for 10min at RT. To quench the crosslinking reaction, cells were incubated 15 min with 0.1 mM Glycine. Unbound particles were then gently washed away 3x with 0.1M DPBS before conducting the staining steps.

As pretreatment for the staining, slides were placed 1 min in ice cold 1% Triton-X100-PBS, washed 3x with PBS and then blocked with 1% BSA-PBS at RT for 30 min. Cells were then stained with DAPI (5mg/mL, 1:5000) and Phalloidin (1:200), before being mounted with anti-fade mounting medium and then dried for 2h at RT. Slides were imaged with a SP8 Leica confocal using a 60x objective. Image representation and intracellular dot-based analysis was done with ImageJ.

### *In vivo* experiments

Female Balb/c mice were obtained from Charles River Laboratories. The animals were housed under a 12-h light/dark cycle and had unlimited access to food and water. The temperature was maintained at 18–23 °C, and humidity was at 40–60%. The animals were housed for at least 1 week before entering any experiments. The animal experiments were conducted according to institutional guidelines of the German Mouse Facility at Helmholtz Munich and after approval of the Ethical Review Board of the Government of Upper Bavaria (Regierung von Oberbayern, Munich, Germany) and under European Directive 2010/63/EU for animal research.

### Tissue clearing of animals (nanoDISCO)

All experiments were performed in triplicates. Mice were deeply anesthetized using a combination of ketamine (70mg/kg) and xylazine (20mg/kg) intraperitoneally. Afterwards, the chest cavity of mice was opened for intracardial perfusion with cold heparinized PBS (10–25 U ml−1 heparin dissolved in 0.01 M PBS) using 100–125-mmHg pressure for 5–10 min at room temperature until the blood was washed out. Next, the mice were perfused with cold 4% paraformaldehyde (PFA) to fix the entire mouse body until stiffness was observed in the tail of the mice. Finally, the skin was carefully removed, and the bodies were fixed in 4% PFA overnight at 4 °C and then transferred to PBS for further processing or directly cleared.

The mice were rendered transparent using the nanoDISCO clearing method previously described (Ref Roger paper). In short, mice were placed in a glass chamber and set up in a transcardial circulatory system using peristaltic pumps. After fixation (see above), the mice were intracardially perfused with PBS 3x for 3h, then 2x for 12h with a decolorization solution (1:4 dilution of CUBIC reagent, consisting of 25 wt% N,N,N,N′-tetrakis (2-hydroxypropyl) ethylenediamine (Sigma-Aldrich, 122262) and 15 wt% Triton X-100 (Fisher Scientific, 10287923)). After this, the mouse bodies were washed with PBS for 3×4h, and then perfused 2d with a decalcification solution (10 wt/vol% EDTA, pH 8.00). For visualizing a direct signal (such as AF-labeled mRNA), no additional staining was needed and the clearing protocol continued as follows: After another three PBS washes lasting 4h each, the mice were transferred to passive clearing (no transcardial pumping) on a shaker under a hood. We proceeded to incubate the mice in 70 vol% THF (Carl Roth, CP82.1), 90 vol% THF and 2 × 100 vol% THF. This was followed by a 20-min treatment with DCM (KK47.1, Carl Roth) and finally refractive index matching using BABB (2:1 solution containing Benzyl Benzoate (Sigma-Aldrich, W213802) and Benzyl Alcohol(Sigma-Aldrich, 1009812500) until the mouse bodies were transparent.

For visualizing an mRNA-transcribed EGFP signal, the protocol after the decalcification step continues as follows: After another three PBS washes lasting 4h each, mice were placed in a permeabilization solution containing 35 ul 0.5 g/L signal-enhancing nanobodies(Alexa Fluor 647–conjugated anti-GFP signal-enhancing nanobodies, ChromoTek, gb2AF647). The permeabilization solution (0.5% Triton X-100, 1.5% goat serum (Gibco, 16210072), 0.5 mM methyl-beta-cyclodextrin (Sigma-Aldrich, 332615) and 0.2% trans-1-acetyl-4-hydroxy-L-proline (441562, Sigma-Aldrich) was actively pumped transcardially for 5 d at RT and then passively incubated with the mice for an additional 2d. Washing of unbound nanobodies was done with a washing solution (containing 0.5% Triton X-100 and 1.5% goat serum) that was applied 2×12h and 3×4h PBS wash steps. As described above, mice were then passively cleared at room temperature with gentle shaking under a fume hood by incubation in 70 vol% THF, 90 vol% THF, 100 vol% THF and again 100 vol% THF for 12 h each. Then, the mouse bodies were treated with DCM for 20 min and immersed BABB until the tissue was rendered completely transparent.

### Flow cytometry (FACS)-based immune cell type analysis in animals

All experiments were performed with n=5. 1µg mRNA containing particles (LNPs or LipiGo particles) were intravenously injected into wildtype Balb/c mice and circulated for 1h, before sacrificing the mice via Ketamin/Xylazin overdose, administered intraperitoneally. After administering the anaesthesia, antibodies were injected 3 min prior to perfusion to label intravascular CD45+ cells. Before being perfused with 0.1M PBS, blood was taken from the hearts of the mice from which serum was generated for further analysis. After the perfusion, organs (spleen, liver, lymph node, bone marrow (femur)) were taken from the mice and placed directly into PBS on ice. Mononuclear phagocytes were isolated by enzymatic digestion and density gradient centrifugation, adapted from the reference^27^.

For processing the liver, organs were transferred into petri dishes for mechanical dissociation with syringe backs until homogenized. The resulting tissue slurry was incubated in digestion buffer containing PBS supplemented with 450 U/mL collagenase I, 125 U/mL collagenase XI, 60 U/mL DNase I, and 60 U/mL hyaluronidase I-S (all from Sigma-Aldrich) at 37 °C for 30 min under agitation. Digested tissues were filtered through a 40 µm cell strainer into 50 mL tubes, followed by centrifugation at 500 g for 7 min at 4 °C. The resulting cell pellets were resuspended in 5 mL of 1× PBS and layered over 5 mL of Histopaque-1077 (Sigma-Aldrich) at room temperature. Samples were centrifuged at 300 g for 20 min (acceleration 3, brake 0), and the mononuclear cell layer at the interphase was collected and washed in FACS buffer (Cell Staining Buffer (420201, BioLegend)).

Lymph nodes and the spleen were treated with a slightly altered protocol, where the tissue was mechanically dissociated through a 40-µm cell strainer into 20 mL of ice-cold PBS. The cell suspension was pelleted (500 × g, 7 min, 4 °C), washed once with 1 mL of FACS buffer, and centrifuged again. Bone marrow was harvested by removing the femoral epiphyses and flushing the marrow cavity with ice-cold PBS using a 23-gauge needle. The cell suspension was passed through a 40-µm cell strainer, and the cells were pelleted (500 × g, 7 min, 4 °C).

After these pretreatments, a common staining step followed, where isolated cells were centrifuged at 500 g for 7 min at 4 °C and resuspended in 50–80 µL of FACS buffer (Cell Staining Buffer, 420201, BioLegend). To block Fc receptors, 50 µL of buffer containing 0.5 µL Fc-block (CD16/CD32 Monoclonal Antibody (93), eBioscience™, 14-0161-85, Thermo Fischer) and 0.25 µL Zombie NIR viability dye (BioLegend) was added, followed by incubation for 10 min at RT°C. Subsequently, 50 µL of antibody cocktail was added per tube (antibodies listed below), and samples were incubated for 30 min at 4 °C. Cells were washed with 1 mL FACS buffer and centrifuged again at 500 g for 7 min. Final pellets were resuspended in 250 µL of FACS buffer for acquisition. The cells were measured by a Cytek® Northern Lights™ 3000.

The following fluorochrome-conjugated antibodies were used for surface staining: CD3 (clone 17A2, BioLegend), CD45 i.v. (clone 30-F11), CD8a (clone 53-6.7, BD Biosciences), CD11b (clone M1/70, BioLegend), CD35 (clone 8C12, BD Biosciences), CD62L (clone MEL-14, BD Biosciences), CD5 (clone 53-7.3, BD Biosciences), B220 (clone RA3-6B2, BioLegend), MHC-II (clone M5/114.15.2, Invitrogen), CD44 (clone IM7, Invitrogen), Ly6G (clone 1A8-Ly6g, Invitrogen), Ly6C (clone AL-21, BD Biosciences), F4/80 (clone BM8, Invitrogen), CD11c (clone N418, Invitrogen), CD45 (clone 30-F11, BioLegend), and CD4 (clone GK1.5, BioLegend). Cell viability was determined using ZombieNIR dye (BioLegend). TX45), anti-MHC-II (FITC, clone M5/114.15.2), anti-Ly6C (PE-eFluor610, clone 1A8-Ly6g), anti-Ly6G (PerCP/Cy5.5, clone HK1.4), anti-F4/80 (PE/Cy7, clone BM8), anti-SIRPα (AlexaFluor700, clone P84), and anti-CD45 (APC/Cy7, clone 30-F11). The antibody mix was generated in the buffer BD Horizon™ Brilliant Stain Buffer (566349, BD Biosciences).

### Bioluminescence imaging

All experiments were performed in triplicates. Mice were injected with LNPs/LipiGo particles encapsulating FLuc-mRNA 24h before imaging. 10 min before imaging the mice were deeply anaesthezized with a Ketamin/Xylazin solution and injected with 150ul of a 15mg/ml aqueous Luciferin solution (VivoGlo luciferin, Promega). intraperitoneally. The animals were then scanned on an IVIS bioluminescent scanner as a whole, before being terminated via cervical dislocation. The organs from the sacrificed mice were then quickly collected for an additional, out-of-body scanning. The exposure time for each scan was set to 60s. The quantifications were assessed of each organ by marking them as individual ROIs to generate the desired values using the Living Image Software version 4.2 (Caliper Life Sciences).

### Multiphoton imaging for particle Half-Life in blood assessments

Multiphoton imaging was performed using an upright Zeiss LSM710 confocal microscope equipped with a Ti:Sa laser (Coherent, Chameleon Vision II) and two external photomultiplier detectors for red and green fluorescence channels. Eight-week-old C57BL/6N mice (Charles River Laboratories) were anesthetized via intraperitoneal injection of a medetomidine (0.5 mg kg⁻¹), fentanyl (0.05 mg kg⁻¹), and midazolam (5 mg kg⁻¹) mixture (MMF). Body temperature was continuously monitored and maintained using a rectal probe connected to a feedback-controlled heating pad.

A catheter was inserted into the femoral artery for administration of fluorescent dyes or nanoparticles. A 4 × 4-mm cranial window was surgically prepared over the right frontoparietal cortex under continuous saline cooling, as previously described. To visualize brain vasculature and acquire baseline images, mice were injected intravenously with FITC-dextran (3 μl g⁻¹). The animal was then positioned on the multiphoton microscope platform, which was adapted for intravital imaging of small animals.

Mice was anesthetized, and a rectangular 4×4-mm cranial window was drilled over the right frontoparietal cortex under continuous cooling with saline. After implantation of glass cranial window, mouse was placed under the microscope and the brain vessels were imaged to obtain a baseline. Afterwards, the nanoparticles labeled with 0.5 mol % FastDIO (0.067ug/kg in a volume of 100 μl.) were systemically injected, and serial imaging was immediately started for at least 2 h. Time-lapse imaging was conducted at an 80-μm tissue depth using 10% laser power at 800 nm. For the FITC channel, a GAASP detector with a long-pass (<570 nm) filter and master gain of 600 was used. The particle channel was recorded using a long-pass (>570 nm) filter and the same master gain setting. Fluorescence signal analysis was performed using ImageJ.

### Light-sheet imaging and image visualization of whole cleared mice

Whole-body imaging of cleared mice was performed using a light-sheet fluorescence microscope (Miltenyi Blaze Ultra) equipped with a 4× objective. Mice were mounted ventrally and a z-step size of 6 µm was applied. In the juvenile mice used in this study (<6 weeks of age), a Z-range of 1200–1400 µm was typically sufficient to capture the entire body. To improve sensitivity for detecting nanoparticles, the light-sheet width was reduced to 80%. Scanning was conducted in “light speed mode,” enabling rapid acquisition.

Laser settings were adjusted based on fluorophore characteristics: for 647-labelled mRNA, laser power was set to 100% with an exposure time of 15 s; for background imaging, 80% power and a 5 ms exposure were used. Prior to full dataset transfer, a subset of image tiles was stitched and assessed for image quality to ensure suitability for downstream analysis. Tile stitching was performed on high-performance workstations using a custom Python script. Two-dimensional image stacks were converted to 3D volumes using Imaris Converter, and image visualization, including snapshot generation, was carried out in Imaris Viewer.

### SCP-nano particle quantification

The in-house developed SCP-nano pipeline consists of organ segmentation, optional manual organ segmentation correction using VR, and subsequent blob detection, and was described in detail in Luo et al.^15^. In short, for quantifying the particle events per organ, the relative intensity contrast of every segmented LNP dot compared to the background intensity was calculated. The background intensity was estimated per organ by the average intensity of voxels inside the organ while excluding voxels belonging to segmented dots. The relative contrast values of all segmented dots were summed up to reflect the amount of LNP in a specific area or organ. Density heat maps could then be generated based on nanoparticle segmentation to visualize nanoparticle distribution in different organs. A sliding window strategy was employed to compute segmented nanoparticles’ local relative intensity contrast values. A window size of 16 × 16 × 4 voxels was deployed in our experiments. Every voxel in this cube was then assigned with the sum of relative contrast values. Finally, using a 3D Gaussian filter, we obtained a smooth and full-resolution density map.

### Immunostaining of tissues and slide scanner imaging

Fixed tissue pieces of perfused mice were dehydrated in a 30% sucrose solution and then sliced in 10 µm slices with a Cryostat and collected on glass slides. 6 slices in per sample were then stained with a classical immunostaining protocol, consisting of 3x 5 min PBS wash, a 15 min permeabilization step with 0.1% Triton-X100, a 1h blocking step with 3% BSA and an overnight staining step at 4°C with a primary antibody in a buffer containing 0.1% Triton and 3% BSA. Unbound antibody was removed by washing 3x with PBS, before staining with a secondary antibody for 2h at room temperature. Samples were additionally stained with Hoechst (3 min with 7 µM), before being washed 3x with PBS and 1x with ddH_2_O. After drying, the mounting medium (VECTASHIELD Antifade, VEC-H-1000, Biozol) was used and slides were closed with cover slips, before being scanned with a slide scanner at the Helmholtz Imaging Core Facility. Automated image analysis was done using the Visiopharm software and plots generated with GraphPad Prism.

## Acknowledgements

This work was supported by the European Research Council Consolidator grant (no. GA 865323); Nomis Heart Atlas project grant (Nomis Foundation); Helmholtz Association Program Helmholtz IVF (ZT-I-PF-4-091, FOMIA); and the German Research Foundation (DFG) under Germany’s Excellence Strategy (EXC 2145 SyNergy – ID 390857198). S.H. acknowledges funding from the Amgen Scholarship program.

We thank Helmholtz Core Facility for Electron Microscopy (Dr. Carsten Peters) for guidance with cryo-TEM imaging, and the Helmholtz Core Facility Imaging (Dr. Annette Feuchtinger) for support with slide scanner imaging and analysis. We also thank Prof. J. Cui for access to dynamic light scattering measurements, Prof. Ernst Wagner for providing HeLa-Gal8-mRuby3 cells, Dr. David Jürgens for assistance with cytokine assays, and Dr. Doris Kaltenecker for serum protein analysis.

## Contributions

A.E. conceived the project as a whole, including the sections on particle design, imaging, and AI analysis. K.K. conducted the particle design, and K.K. and C.K. conducted majority of the experiments involving particle production, characterization, tissue clearing, light-sheet imaging and data analysis. S.U. and V.K.R. designed and V.K.R. conducted the adipocyte cell culture experiments. A.L conceptualized, and O.C. and A.C. conducted the experiments on immune cell analysis with FACS. D.J. conducted all i.v injections, and assisted with the biomuniescence, tissue clearing, and flow cytometry experiments. K.K. and S.H. planned and S.H. executed experiments involving aptamer decoration of particles. I.K. and N.P. planned and I.K. executed experiments involving intravitreal imaging. Y.C and I.H. worked on the organ segmentation and LNP quantification, L.H. and E.A. conducted corrections using VR tools. D.-P.M. performed proteomics sample preparation and proteomics analysis, and M.A. conducted the proteomics data analysis. T.S. and E.S. helped set up and analyze experiments on single particle profiling of the particles. Y.A.K. contributed to tissue clearing and imaging. J.G. performed immunohistology analysis of the particles in the tissues. A.E., M.E., F.H., and C.K. provided overall supervision for the project. K.K. and C.K. wrote the manuscript; C.K., F.H, and M.E. revised the text.

