## Supplemental Information for "LipiGo: A Versatile DNA-Lipid Nanoparticle Hybrid for Precision Drug Delivery"

### SI Figure 1:Characterization of biophysical properties of particles

a  DLS data of std LNP and LipiGo formulations with different ionizable lipids

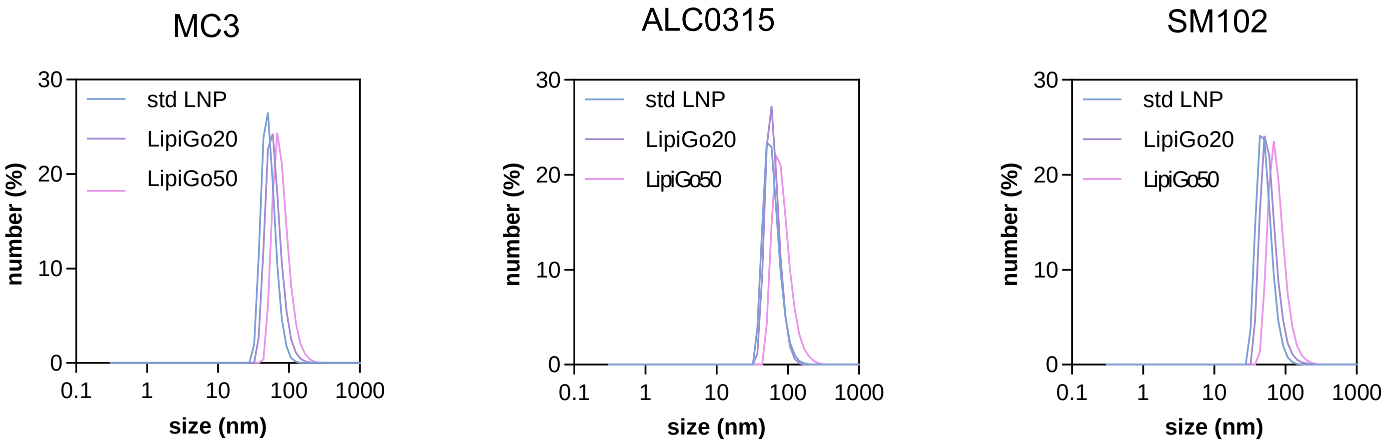


b  DLS data of std LNP and LipiGo formulations with different DNA handles

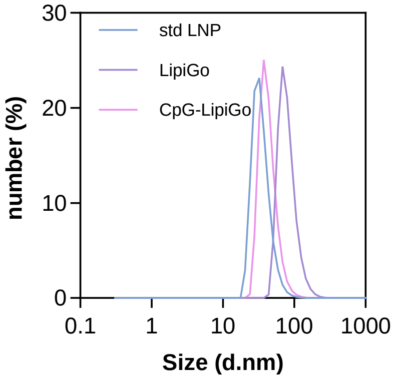


LipiGo+Aptamer characterization

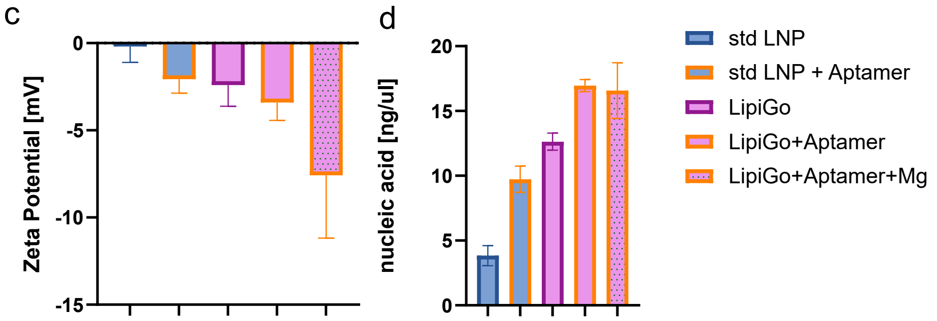


**SI Fig. 1: Characterization of biophysical properties of particles**. **a.** DLS data comparing different lipid base formulations, featuring MC3, ALC0315 and SM102. Shown as mean, n=3 **b.** DLS data comparing different handle sequences, featuring a random sequence and a CpG motiv sequence of the same length. Shown as mean, n=3. **c.** Zeta potential of aptamer-decorated particles, comparing std LNP, std LNP+Aptamer, LipiGo, LipiGo+Aptamer and LipiGo+Aptamer+Magnesium buffer. Shown as mean with SD, n=3. **d.** nucleic acid content comparing the aptamer-decorated particles listed in c. Measured spectroscopically (nanodrop) and shown as mean with SD, n=3.

### SI Figure 2: Immunogenicity and Toxicity profiling of LipiGo

#### a WST cell viability assay

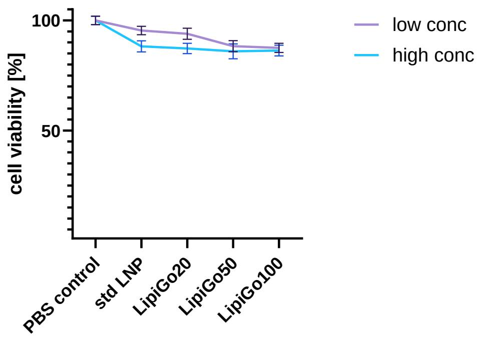


#### b Flow Cytometry-based Cytokine panel

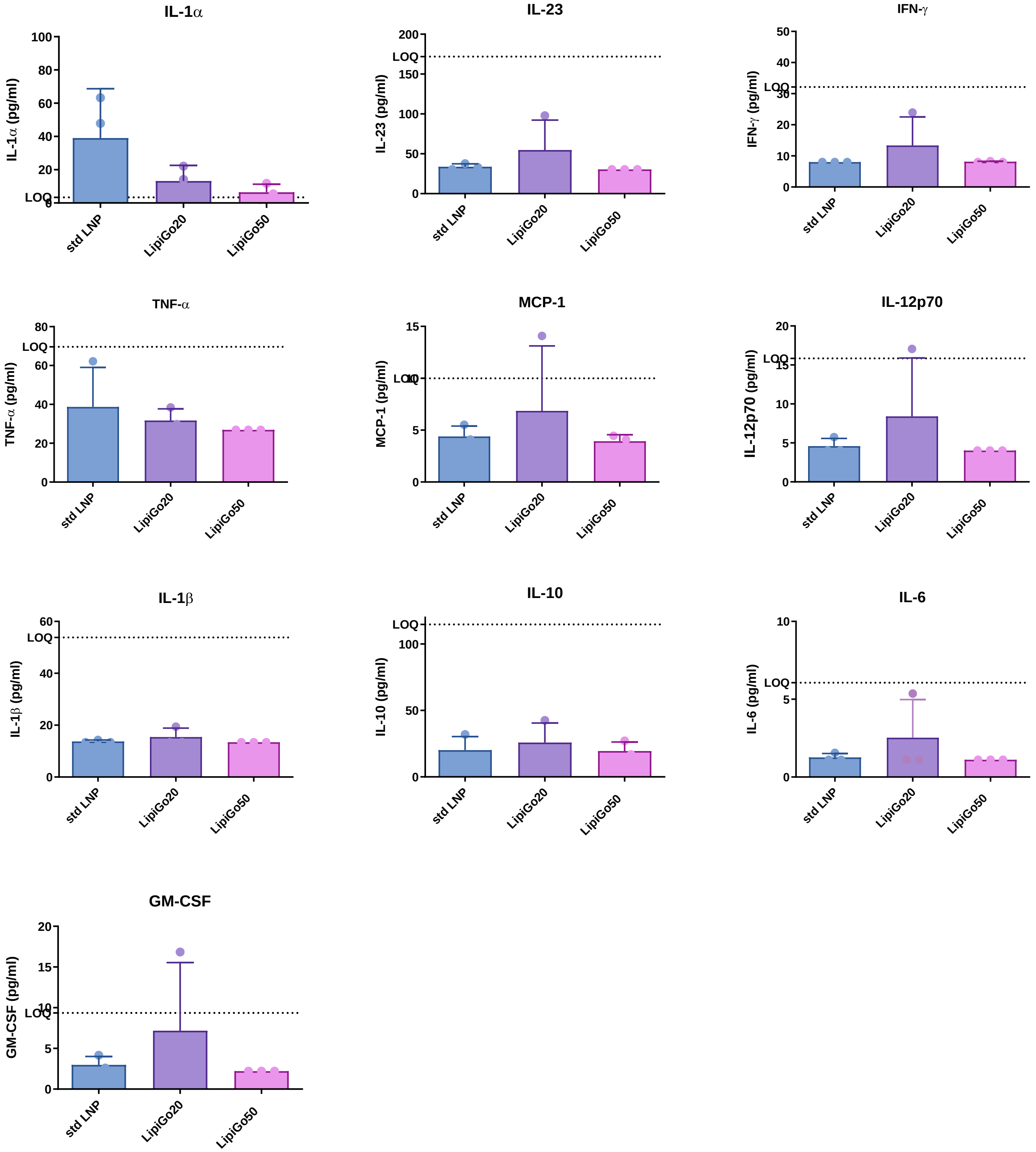

c Serum protein analysis

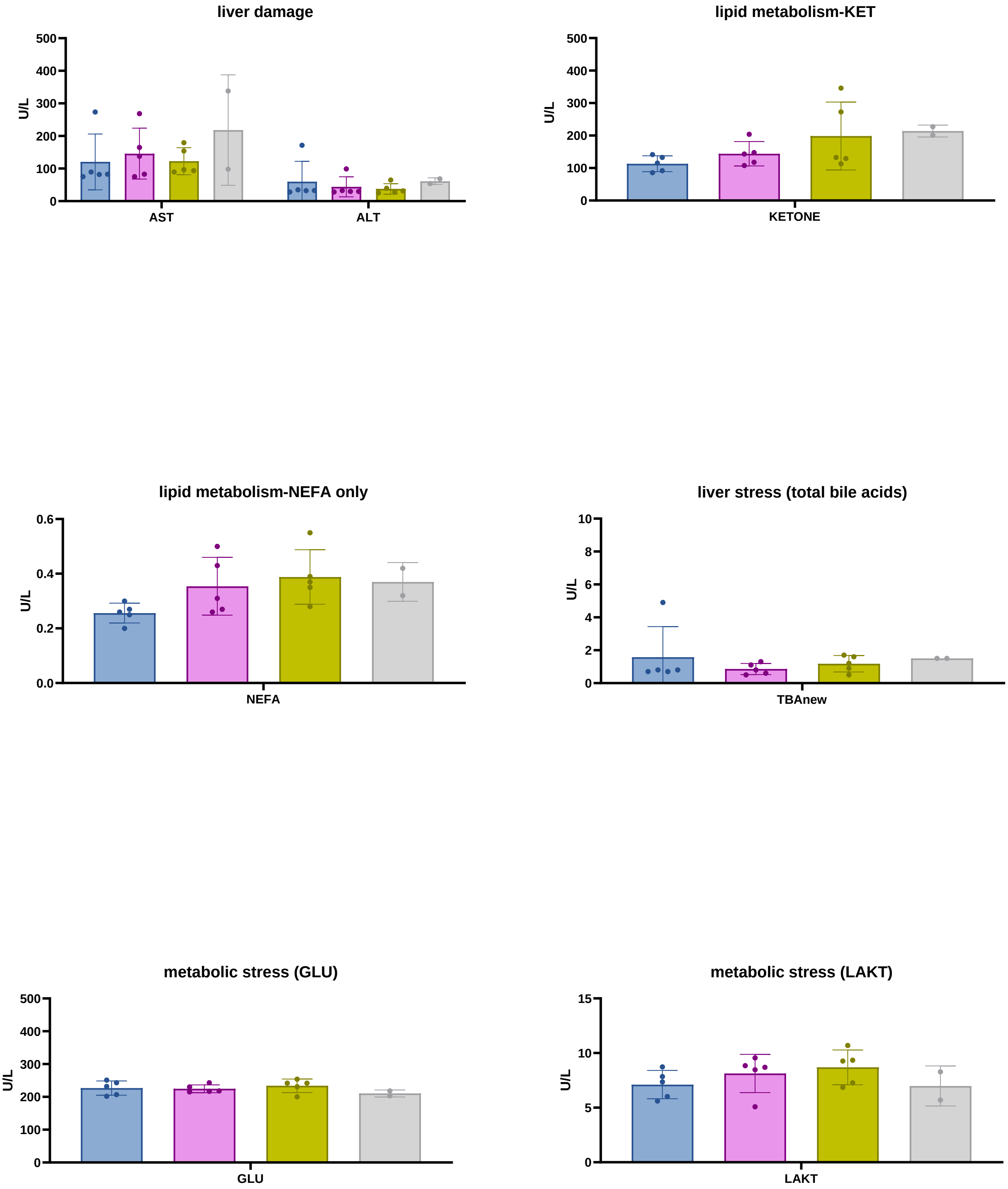


**SI Fig.2:Immunogenicity and Toxicity profiling of LipiGo a. HEK cell based cell viability assay (WST).** n=3, mean and SD shown for 2 different concentrations. **b. 13-plex Immunological panel**, measured from the serum of animals 48h after injection with 2.4ug mRNA packaged into std LNP (blue), LipiGo20(purple) or LipiGo50(pink). **c. Analysis of serum proteins.** After 1h of i.v. injection of 10ug mRNA packaged in standard LNP (blue), LipiGo50(pink) or LipiGo50-CpG(green). Negative control PBS injected (grey)

### SI Figure 3: Biodistribution of particles in i.v. injected animals.

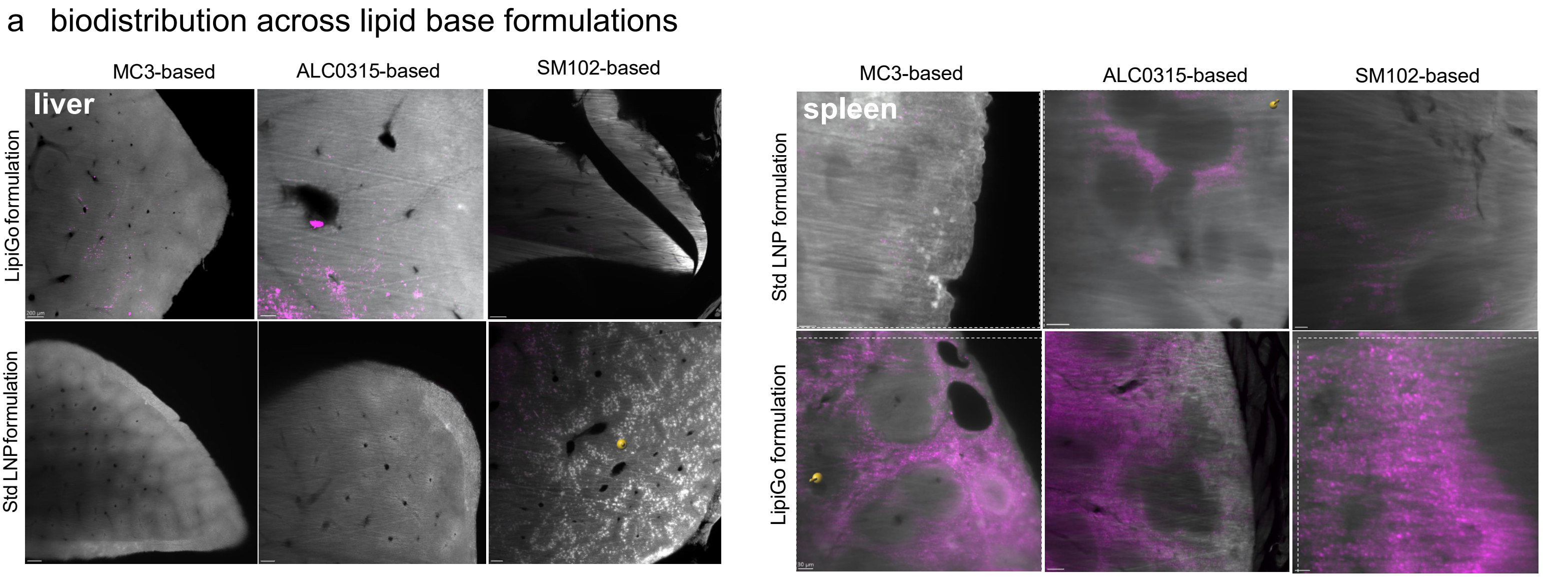


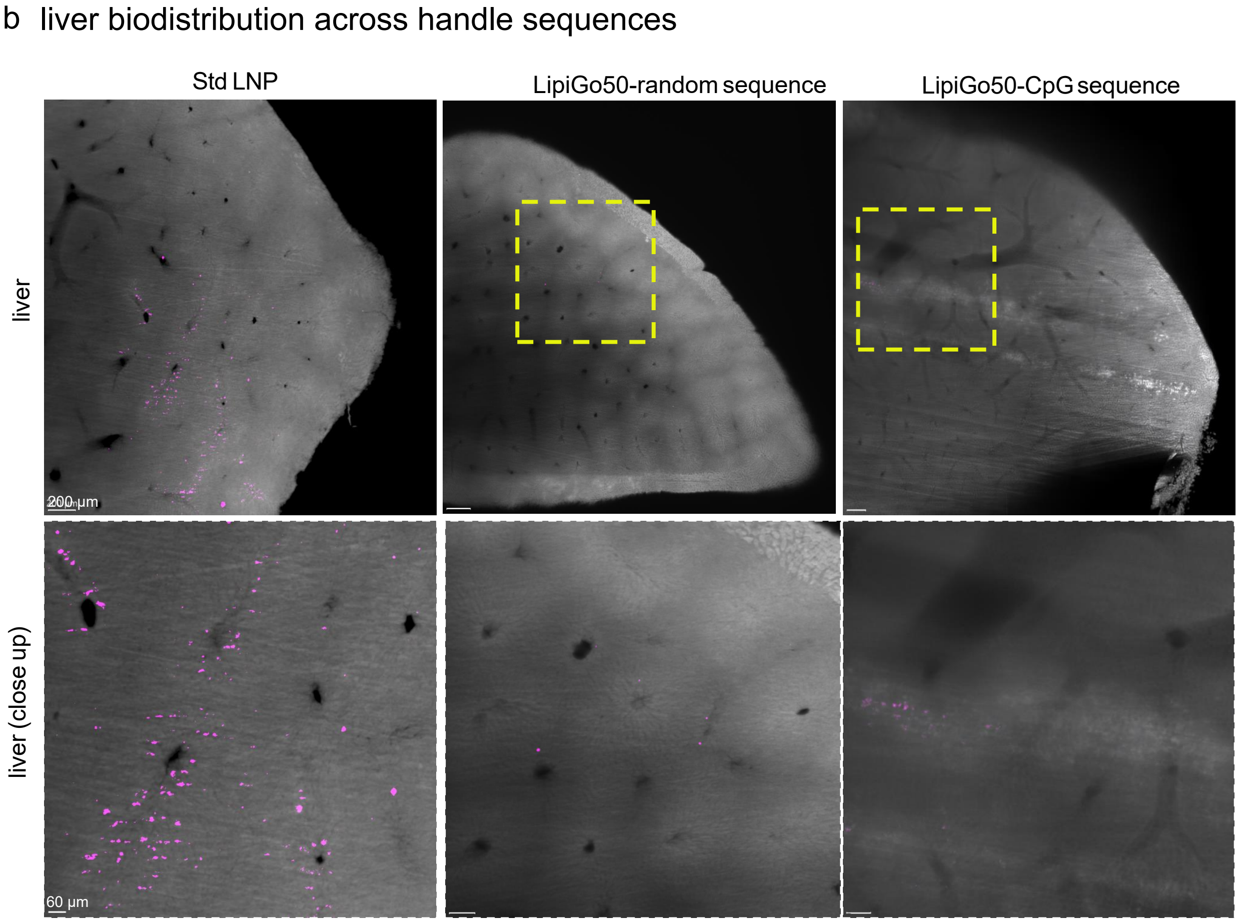

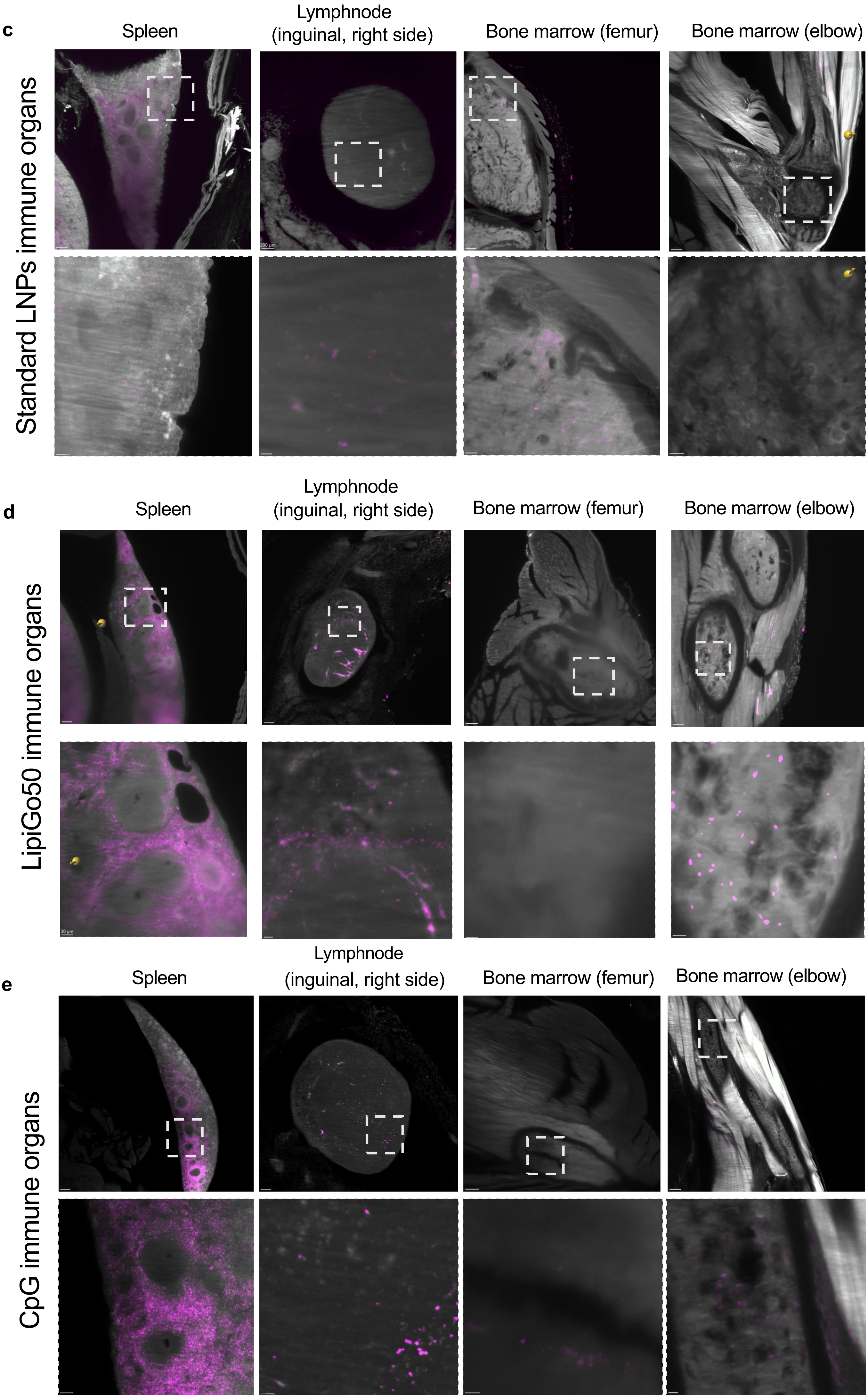
c
Spleen
Lymphnode
(inguinal, right side)
Bone marrow (femur)
Bone marrow (elbow)
Standard LNPs immune organs
d
Spleen
Lymphnode
(inguinal, right side)
Bone marrow (femur)
Bone marrow (elbow)
LipiGo50 immune organs
e
Spleen
Lymphnode
(inguinal, right side)
Bone marrow (femur)
Bone marrow (elbow)
CpG immune organs

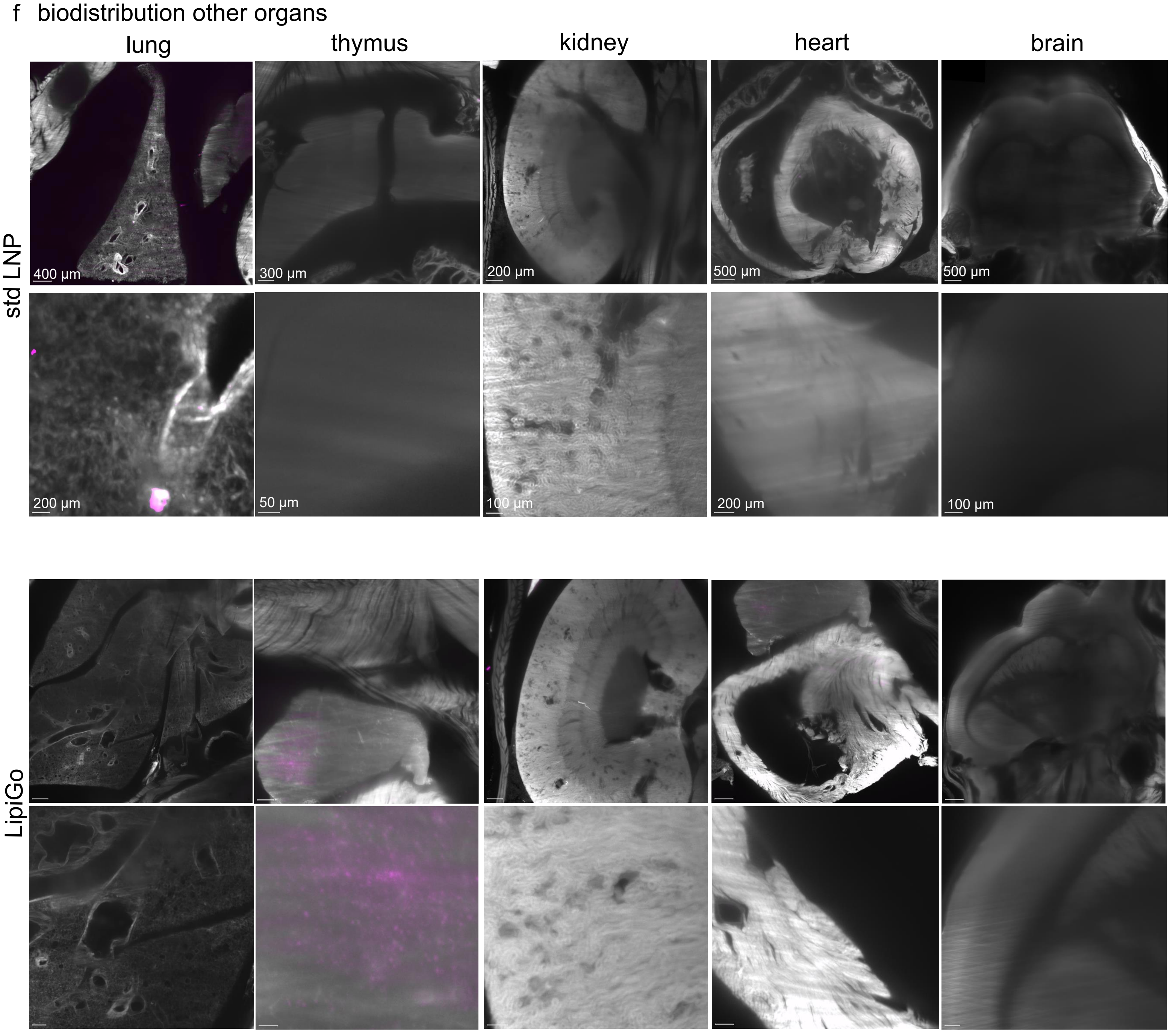


**SI Fig.3: Biodistribution of particles in i.v. injected animals.** Representative light-sheet images of tissue cleared mice that were injected with 1 ug mRNA/mouse encapsulated into different carriers for comparison. **a.** Biodistribution to liver and spleen comparing LipiGo and std LNP variants with different lipid base formulations. Particles were either formulated with the ionizable lipid MC3, ALC0315 or SM-102. **b.** Liver biodistribution across different DNA handles, comparing no handle (std LNP), a random DNA sequence and a CpG-motif DNA sequence. **c,d,e.** Representative images of lymphatic tissues (lymphnodes, bone marrow, and spleen) comparing std LNP(c), LipiGo particles (d) and LipiGo-CpG particles (e)

### SI Figure 4: Functional analysis of delivered mRNA cargo.

a Expression of cargo EGFP-mRNA in tissue cleared mouse, std LNP

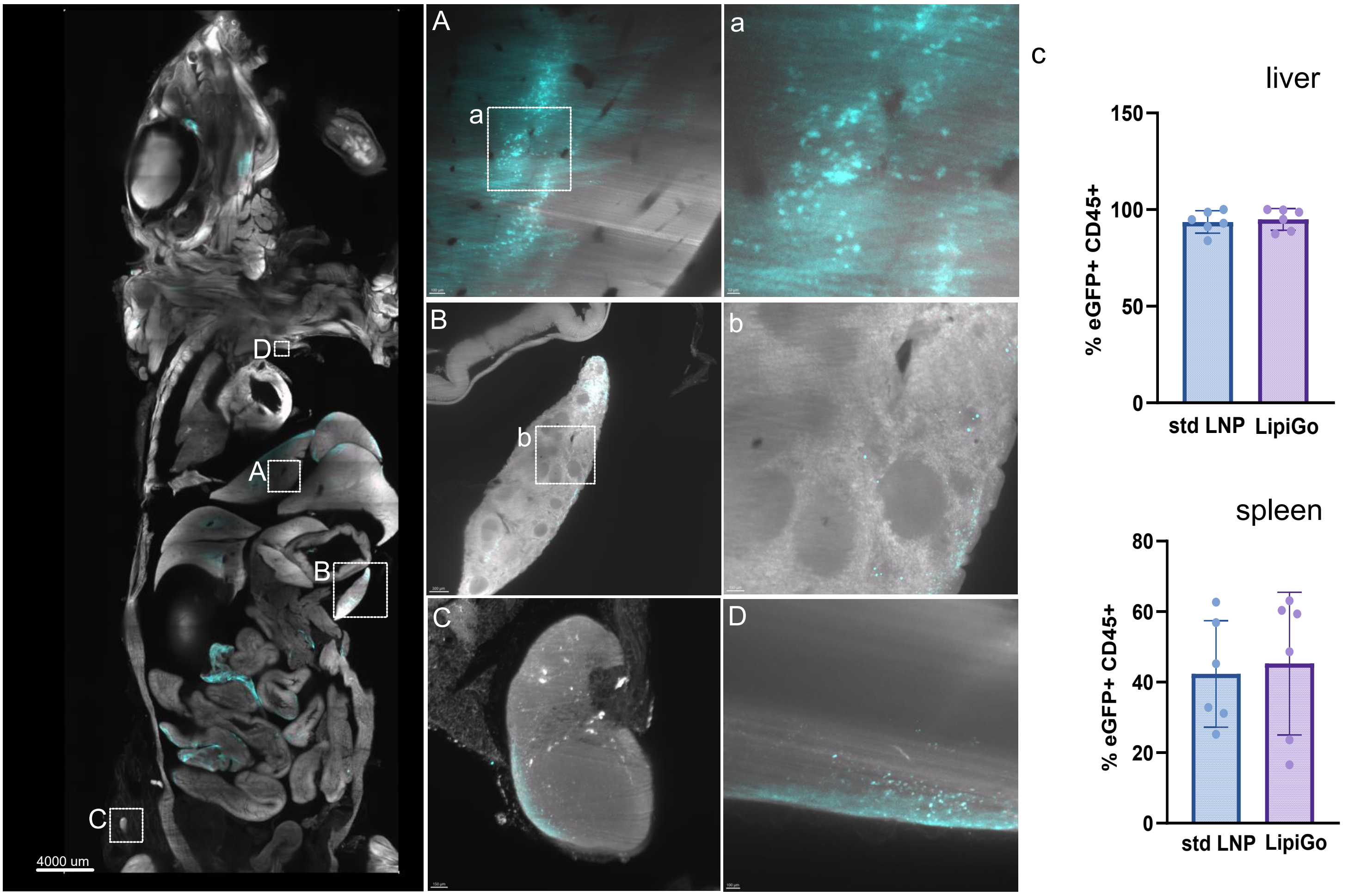


b Bioluminescence data

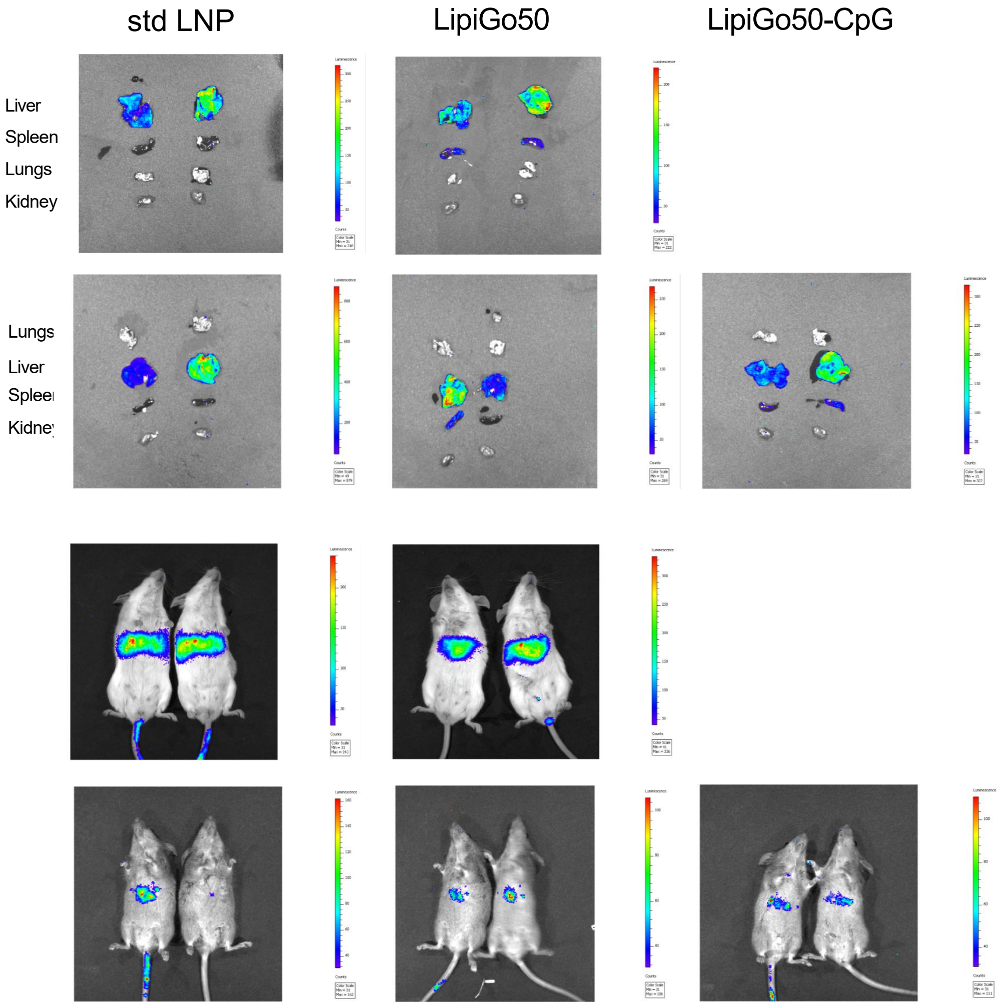


**SI Fig.4: Functional analysis of delivered mRNA cargo. a.** Representative images of std LNP carrying EGFP-mRNA, whole body (left) and organs of interest (right), showing the liver (A), the spleen (B), an inguinal lymphnode (C) and the thymus (D). **b.** Complete bioluminescence data, showing the underlying data of the quantification in main Fig.4. n=4 or 2

### SI Figure 5: Complete Flow Cytometry (FACS) data

a Complete FACS data (in % of all cell types)

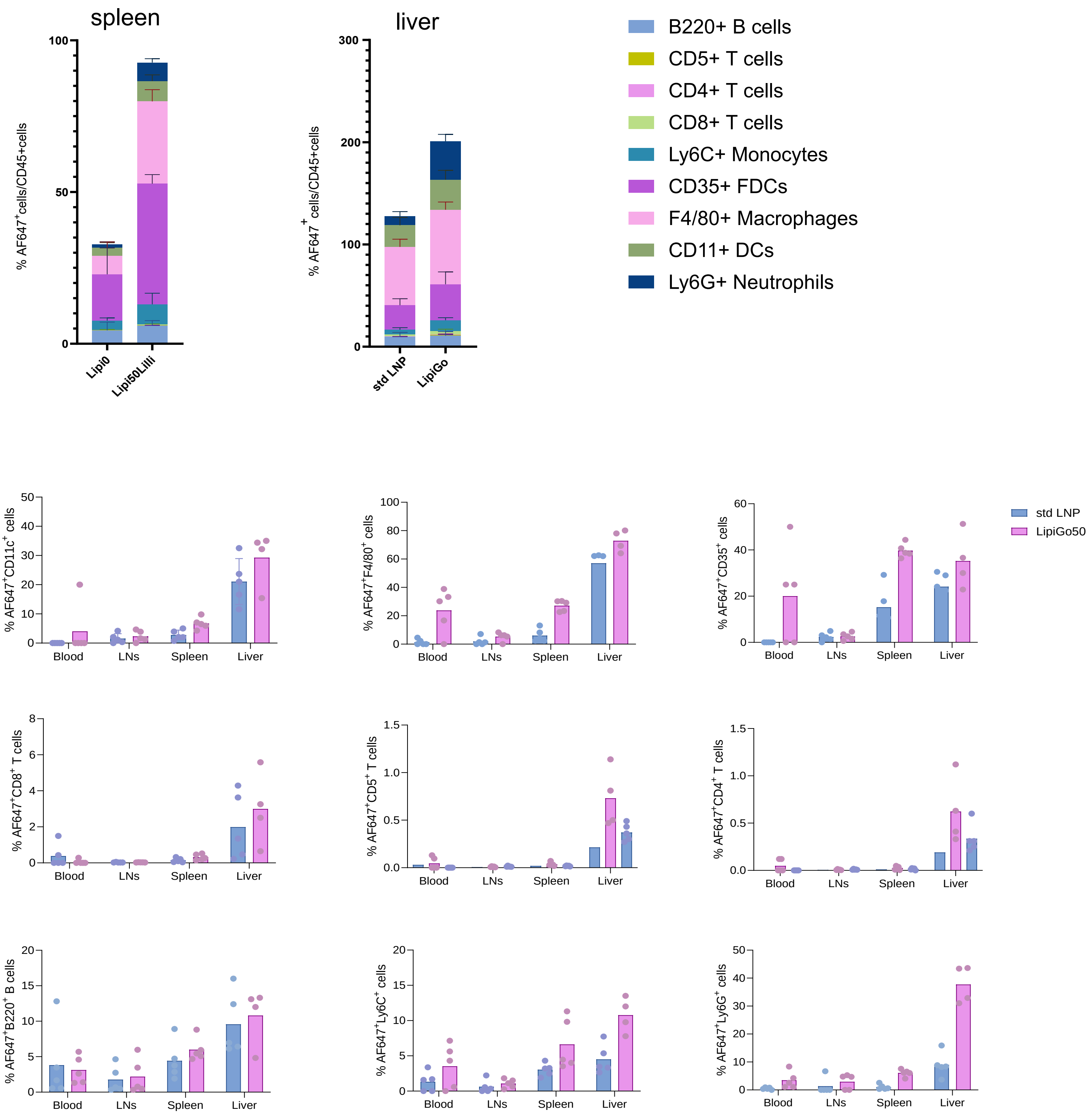

b Complete FACS data (in absolute cell numbers)

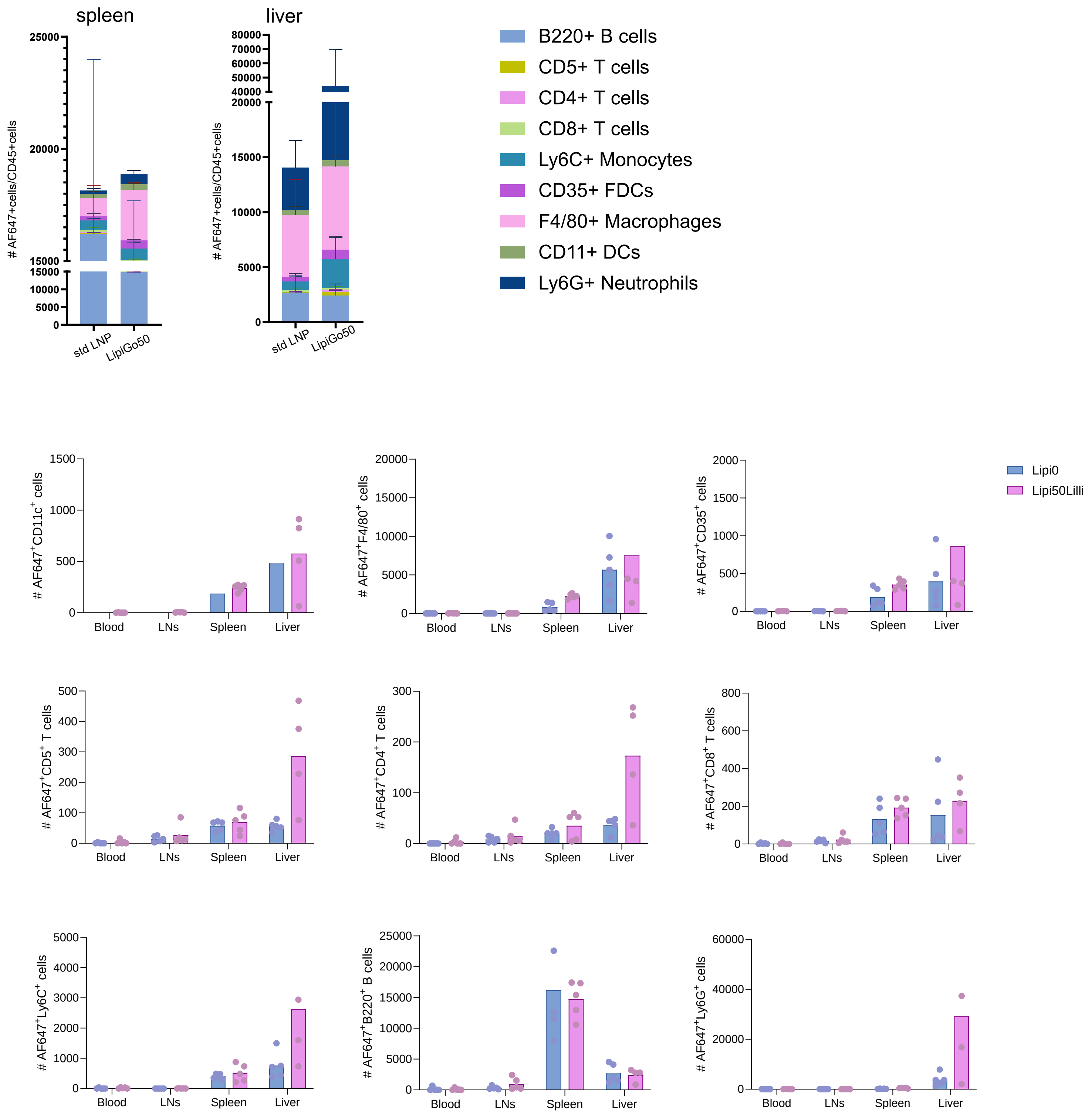


**SI Fig. 5: Complete Flow Cytometry (FACS) data** of fluorescent particles injected intravenously into animals and circulated for 1h. **a**. Data of all measured cell types shown as % of a respective cell type to all cells. **b**. Data of all measured cell types shown as absolute cell numbers of a respective cell type. n=5

### SI Figure 6: Protein corona analysis

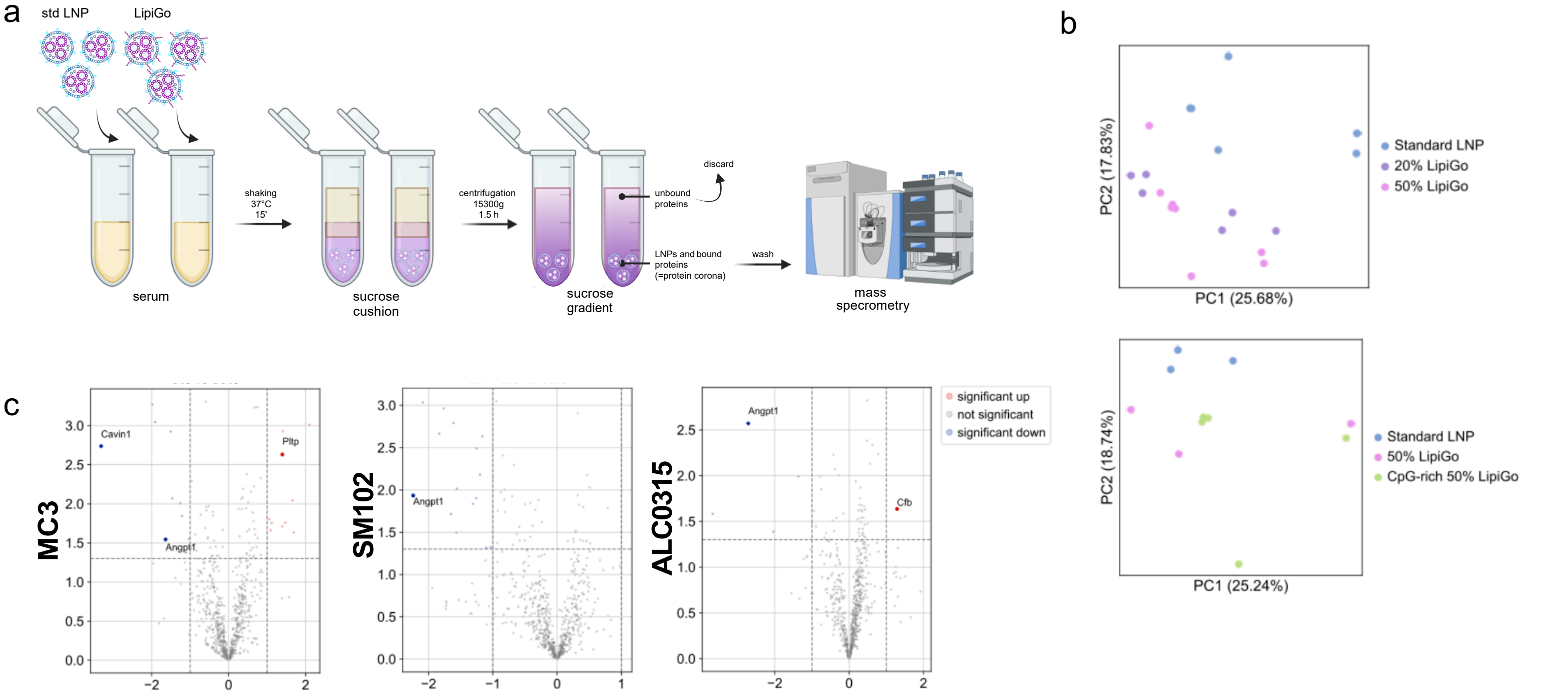


**SI Fig.6 Protein corona analysis a.** schematic of protein corona analysis workflow **b.** PCA plots comparing standard LNP to different LipiGo variations **c.** volcano plots comparing more-and less enriched proteins on the surface of LipiGo variations  with different lipids

#### SI Figure 7: Bioluminescence data aptamer-functionalized LipiGo particles

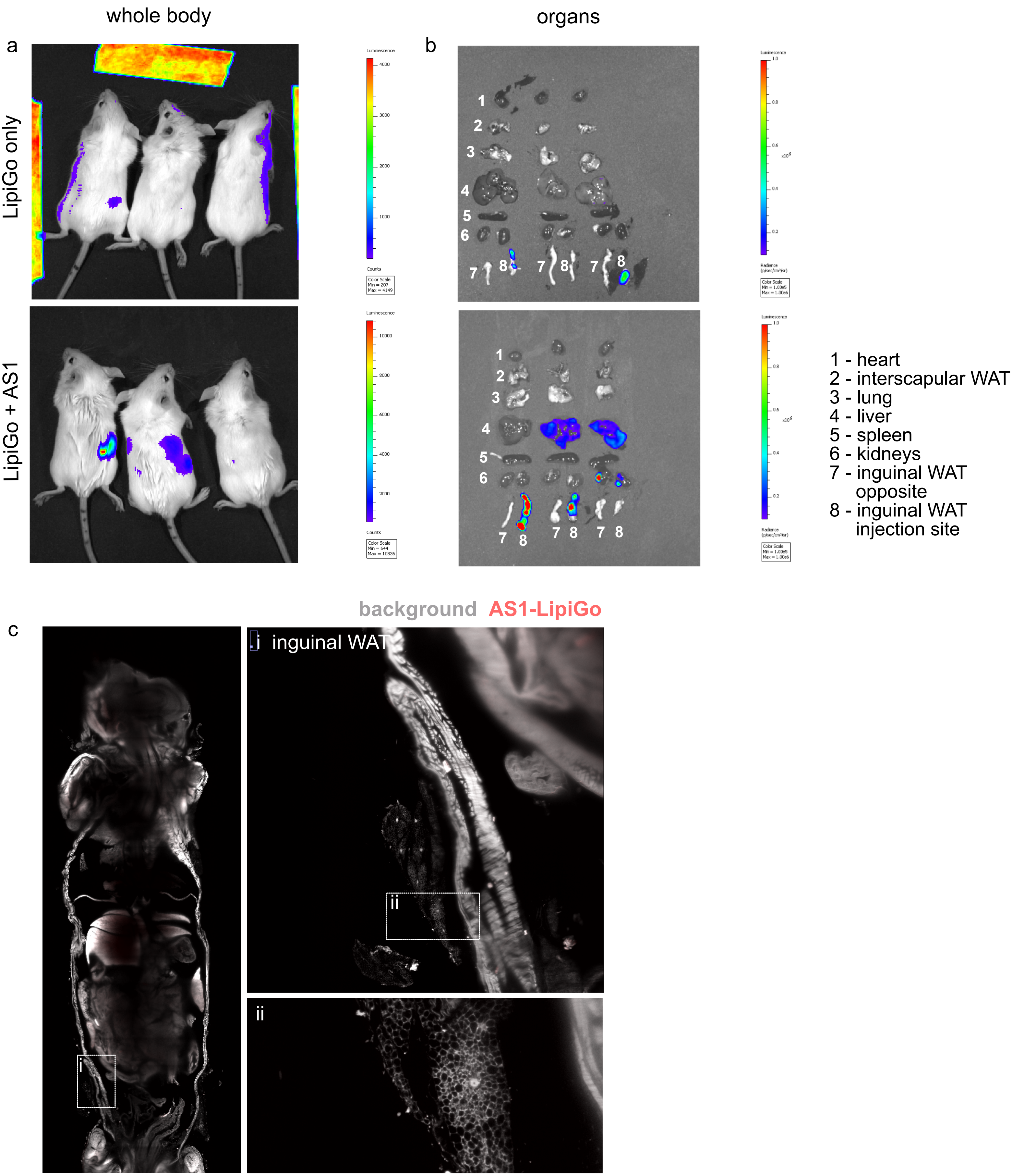


**SI Fig.7 Bioluminescence data aptamer-functionalized LipiGo particles.** 10ug Luciferase mRNA per mouse, n=3 **a.** Whole body images of LipiGo or LipiGo-aptamer injected mice. **b.** Ex vivo organs of the same mice, correlating to the quantification data from main Fig.5. c. Representative image from tissue cleared and light-sheet imaged mouse, injected with LipiGo only. The inguinal WAT close to the iinjection site is highlightes
